# Functional differentiation of Human Dental Pulp Stem Cells into neuron-like cells exhibiting electrophysiological activity

**DOI:** 10.1101/2024.10.18.619039

**Authors:** B. Pardo-Rodríguez, A.M. Baraibar, I. Manero-Roig, J. Luzuriaga, J. Salvador-Moya, Y. Polo, R. Basanta-Torres, F. Unda, S. Mato, G. Ibarretxe, J.R. Pineda

## Abstract

**Background:** Human dental pulp stem cells (hDPSCs) constitute a promising alternative for central nervous system (CNS) cell therapy. Unlike other human stem cells, hDPSCs can be differentiated, without genetic modification, to neural cells that secrete neuroprotective factors. However, a better understanding of their real capacity to give rise to functional neurons and integrate into synaptic networks is still needed. For that, *ex vivo* differentiation protocols must be refined, especially to avoid the use of fetal animal serum.

**Methods:** In this study, we sought to improve existing differentiation protocols for obtaining functional neuron-like cells from hDPSCs. We compared the effects of the absence or presence of fetal serum during the initial expansion phase as a step prior to switching cultures to neurodifferentiation media. We improved hDPSC neurodifferentiation by adding retinoic acid (RA) and potassium chloride (KCl) pulses for 21 or 60 days and characterized the results by immunofluorescence, digital morphometric analysis, RT-qPCR and electrophysiology.

**Results:** We found that neural markers like Nestin, GFAP, S100β and p75^NTR^ were expressed differently in neurodifferentiated hDPSC cultures depending on the presence or absence of serum during the initial cell expansion phase. In addition, hDPSCs previously grown as spheroids in serum-free medium exhibited *in vitro* expression of neuronal markers such as doublecortin (DCX), neuronal nuclear antigen (NeuN), Ankyrin-G and MAP2 after neurodifferentiation. Presynaptic vGLUT2, Synapsin-I, and excitatory glutamatergic and inhibitory GABAergic postsynaptic scaffold proteins and receptor subunits were also present in these neurodifferentiated hDPSCs. Treatment with KCl and RA increased the amount of both voltage-gated Na^+^ and K^+^ channel subunits in neurodifferentiated hDPSCs at the transcript level. Consistently, these cells displayed voltage-dependent K^+^ and TTX-sensitive Na^+^ currents as well as spontaneous electrophysiological activity and repetitive neuronal action potentials with a full baseline potential recovery.

**Conclusion:** Our study demonstrates, for the first time, that hDPSCs can be differentiated to neuronal-like cells that display functional excitability and thus evidence the potential of these easily accessible human stem cells for nerve tissue engineering. Our results highlight the importance of choosing an appropriate culture protocol to successfully neurodifferentiate hDPSCs.

## Background

The central nervous system (CNS) possesses a very limited self-renewal capacity. After a traumatic event or neurodegenerative disease such as Alzheimer’s disease, a progressive loss of synaptic interconnections occurs in neural tissue and patients are left with a chronic and disabling condition that entails considerable dependency burdens [1–4]. Currently, treatments against CNS lesions only relieve symptoms and do not replace damaged neural tissue or disrupt disease progression [5]. Neurogenesis in the adult human brain is rare and endogenous neural stem cells (NSCs) are scarce and difficult to obtain. Hence, in recent years researchers have focused on finding alternative stem cell candidates for neuroregenerative cellular therapies [6].

In this context, human dental pulp stem cells (hDPSCs) can be easily and routinely obtained from adult third molar surgeries. Unlike most stem cells (i.e. embryonic); the use of hDPSCs implies less ethical constraints as they are considered biological waste. Moreover, hDPSCs are able to secrete anti-inflammatory factors [7] and neurotrophins such as brain derived neurotrophic factor (BDNF) or neurotrophin-3 (NT-3) [8,9]. Thus, in the context of brain injury, they are not only good candidates to replace the affected tissue, but also have the potential to contribute to neuroprotection of the damaged region, even to ameliorate the chronic inflammation and oxidative stress responsible of neuronal death after degenerative or traumatic CNS damage.

Interestingly, hDPSCs have their embryonic origin in the neural crest, and are thus promising candidates for neural reconstruction therapy [10]. Due to the ectomesenchymal nature of hDPSCs, they show an outstanding multilineage differentiation capacity [11,12] to differentiate into mesenchymal vasculogenic [13,14], osteogenic and adipogenic cells, among many others [15,16]. In the CNS, hDPSCs can also be reprogrammed into neurogenic and gliogenic neural crest progenitors when they grow as spheroids after they are switched from fetal serum-containing to defined serum-free media [8]. Indeed, previous works also identified the expression of neural markers in hDPSCs cultured under similar serum-free spheroid growth conditions [17]. In addition, neurodifferentiated hDPSCs also possess neurotransmitter and neurotrophin receptors [8] that allow them to respond to CNS signals. However, electrophysiological evidence supporting a neuron-like excitability phenotype of these differentiated stem cells is scarce. The presence of TTX-sensitive voltage-dependent sodium channels in hDPSCs was first described by Arthur et al. [18]. Using a sphere-mediated neurogenic induction method, Gervois et al. and Li et al. subsequently obtained neurodifferentiated hDPSCs that were able to generate AP-like fast depolarizations that reproduced only the rising phase of a neuron AP. However, in these studies hDPSCs were not able to recover their baseline membrane potential after these depolarizations, and no more than one AP-like depolarization could be induced on the same cell [17,19]. Despite these works describing the firing of discrete AP-like depolarizations in hDPSCs [17,19], it remains unclear whether these cells possess the ability to fully differentiate into functional neuron-like cells that can maintain long-term electrical excitability. Furthermore, their capacity to form synaptic connections and integrate into a functional synaptic network, either *in vitro* or *in vivo*, is still largely unknown. .

In the present study, we aimed to optimize hDPSC neurodifferentiation protocols, characterize the reversibility of fetal serum-induced changes when cells are switched from serum-containing to serum-free media, and increase their differentiation rates towards neuronal lineages with retinoic acid (RA) and potassium chloride (KCl). This improved protocol induced neuronal cells derived from hDPSCs to express excitatory and inhibitory synaptic proteins such as Synapsin, Gephyrin and postsynaptic density protein 95 (PSD95), as well as components of the initial axon segment such as Ankyrin G. Importantly, these neurodifferentiated hDPSCs also exhibited spontaneous electrophysiological activity and triggered repetitive action potentials (APs) upon membrane depolarization with full baseline potential recovery, revealing the functionality of neural differentiated hDPSCs *in vitro*. Our work expands current knowledge regarding the capacity of hDPSCs to give rise to neuron-like cells as a first step towards their integration and restoration of neural circuits, making them ideal candidates to counterbalance the reduced synaptic plasticity, loss of synapses and disruption of neural network characteristics of Alzheimer’s disease [20,21].

## Materials and methods

### hDPSCs primary cultures

Human third molars from healthy donors between 18 and 30 years of age were collected from dental surgery waste. Pulp extraction and hDPSCs primary culture were performed following previously reported protocols [13,22]. Briefly, after tooth fracture, the dental pulp was collected and digested with an enzymatic solution of 3 mg/mL collagenase (#17018029, Gibco) and 4 mg/mL dispase (#D4693-1G, Merck) for 1 h at 37°C. After centrifugation, each donor’s cell pellet was resuspended and cultured in parallel following two different culture media. On the one hand, we generated standard plastic adherent cultures on conventional tissue culture-treated flasks (#83.3912.002, Sarstedt) with DMEM (#D5796, Sigma-Aldrich) supplemented with 10% fetal bovine serum (FBS; #SV30160.03, HyClone), 100 U/mL penicillin and 150 mg/mL streptomycin (#11528876, Gibco) to generate adherent cell monolayers. On the other hand, a serum-free culture medium was combined with low binding adhesion surfaces (#3814, Corning), using Neurocult basal media supplemented with human Neurocult proliferation supplement (#05751, Stem Cell Technologies), both at 9:1 ratio, and supplemented with Heparin solution 2 μg/mL (#07980, Stem Cell Technologies), EGF 20 ng/mL, and bFGF 10 ng/mL (Peprotech, London, United Kingdom), 2% B-27 without vitamin A (#12587010, Thermo Fisher), 2 mM GlutaMAX (#11500626, Fischer Scientific), 100 U/mL penicillin and 150 mg/mL streptomycin (#11528876, Gibco) to generate free-floating neurogenic dentospheres.

### Neurogenic differentiation

After three weeks of expansion, 1×10^4^ cells were seeded in 24-well plates coated with 1:100 laminin (#L2020, Sigma-Aldrich) and the culture media were changed for 21-60 days to neural differentiation media composed by human Neurocult basal media supplemented with human Neurocult differentiation supplement (#05752, Stem Cell Technologies), 2% B-27 with vitamin A (#17504044, Thermo Fisher), 2 mM GlutaMAX (#11500626, Fischer Scientific), 100 U/mL penicillin and 150 mg/mL streptomycin (#11528876, Gibco). In some cases, 10 μM retinoic acid (RA) (#554720, Sigma Aldrich.) and one hour pulses every two days of 40 mM potassium chloride (KCl) (#141494, PanReac), starting at seventh day of neuroinduction process, were added to the neural differentiation medium adapting minor modifications of the protocol of Bosch et al. 2004. [23].

### Immunocytochemistry

The cells were fixed with 4% paraformaldehyde (PFA) (#158127-500G, Sigma-Aldrich) at room temperature for 10 minutes and permeabilized by incubation with 1% bovine serum albumin (BSA) (#A9647, Sigma-Aldrich) and 0.3% Triton X-100 (#93443, Sigma-Aldrich) diluted in phosphate buffered saline (PBS; #D5652, Sigma-Aldrich). Then, the primary antibodies were incubated overnight at 4°C in a solution of PBS, 1% BSA and 0.1% Tween-20 (#P1379, Sigma-Aldrich). For the assessment of cell stemness, a human Nestin 1:200 (#MAB1259, Biotechne R&D) primary antibody was used. To assess glial differentiation, primary antibodies against glial fibrillary acidic protein (GFAP) 1:500 (#G9269, Sigma-Aldrich), S100β 1:500 (#20311, Dako) and NGFR (p75^NTR^) 1:250 (#Sc-271708, Santa Cruz Biotechnology) were used. Doublecortin (DCX) 1:300 (#ab113435, Abcam) and neural nuclear protein (NeuN) 1:300 (#ab177487, Abcam) primary antibodies were used to assess the neural phenotype of hDPSCs. The presence of synaptic proteins was measured with antibodies targeting the presynaptic marker Synapsin-I 1:200 (#ab64581, Abcam) and the postsynaptic marker PSD95 1:200 (#ab18258, Abcam). For axon initial segment identification, an anti-Ankyrin-G antibody 1:100 (#43GF8, Thermo Fisher,) was used. hDPSCs differentiation into GABAergic lineage cells was assessed by immunostaining for gamma-aminobutyric acid (GABA) 1:100 (#SAB4200721, Sigma-Aldrich) and glutamic acid decarboxylase 65 (GAD65) 1:100 (#ZRB1091, Sigma-Aldrich). The secondary antibodies Alexa Fluor 488, Alexa Fluor 647 and Alexa Fluor 555 donkey anti-rabbit, anti-mouse, and anti-goat (#A31572, #A32766, #A31570, #A21206, #A21432, Invitrogen) and DAPI (#10116287, Thermo Fisher) were incubated in a solution of PBS, 1% BSA and 0.1% Tween-20 at room temperature for 2 hours. Image acquisition was carried out via an LSM800 Zeiss confocal microscope (Jena, Germany) at 20X and 63X magnifications. For presynaptic vesicle protein Synapsin-I detection, a Zeiss LSM880 Fast Airyscan (Jena, Germany) high-resolution fluorescence microscope was used at 63X magnification.

### Quantitative image analysis

To determine the percentage of positive cells, the expression of each marker was measured in duplicates in samples from three different donors, which were cultivated in parallel for each neurodifferentiation protocol. For each sample, five images from different areas were acquired with the same acquisition settings. Each marker expression was quantified by keeping the same parameters in blinded samples via ImageJ public software (version 1.50e) [24]. Nuclear morphometric analyses were conducted via the NII plugin for ImageJ software following previously described procedures [25]. The nuclear area was measured in a total of 1,200 cells per experimental condition. Cellular morphology was analyzed with 3D-Sholl Neuronstudio software 0.9.92 (Computational Neurobiology and Imaging Center, Mount Sinai School of Medicine, New York) as previously described [26]. In this case, for each experimental condition, 30 cell morphologies from 3 different donors were determined by measuring different parameters, such as i) the average branch length, ii) the number of branches, iii) the dendrite volume or iv) the dendritic area.

### Quantitative real-time PCR (qRT‒PCR)

RNA extraction was performed via an RNAqueous-Micro Kit (#AM1931, Thermo Fisher) following the manufacturer’s instructions. For reverse transcription, an iScript cDNA synthesis kit (#170-8890, Bio-Rad) was used, and qPCR was carried out using the Bio-Rad SYBR Green Supermix (#1725120, Bio-Rad). All the reactions were performed in triplicate, and the relative expression of each gene was calculated via the 2^-ΔΔCt^ method [27]. GAPDH and β-actin were used as housekeeping genes. The primers pairs used in this study are listed in **Table 1**.

**Table 1:**
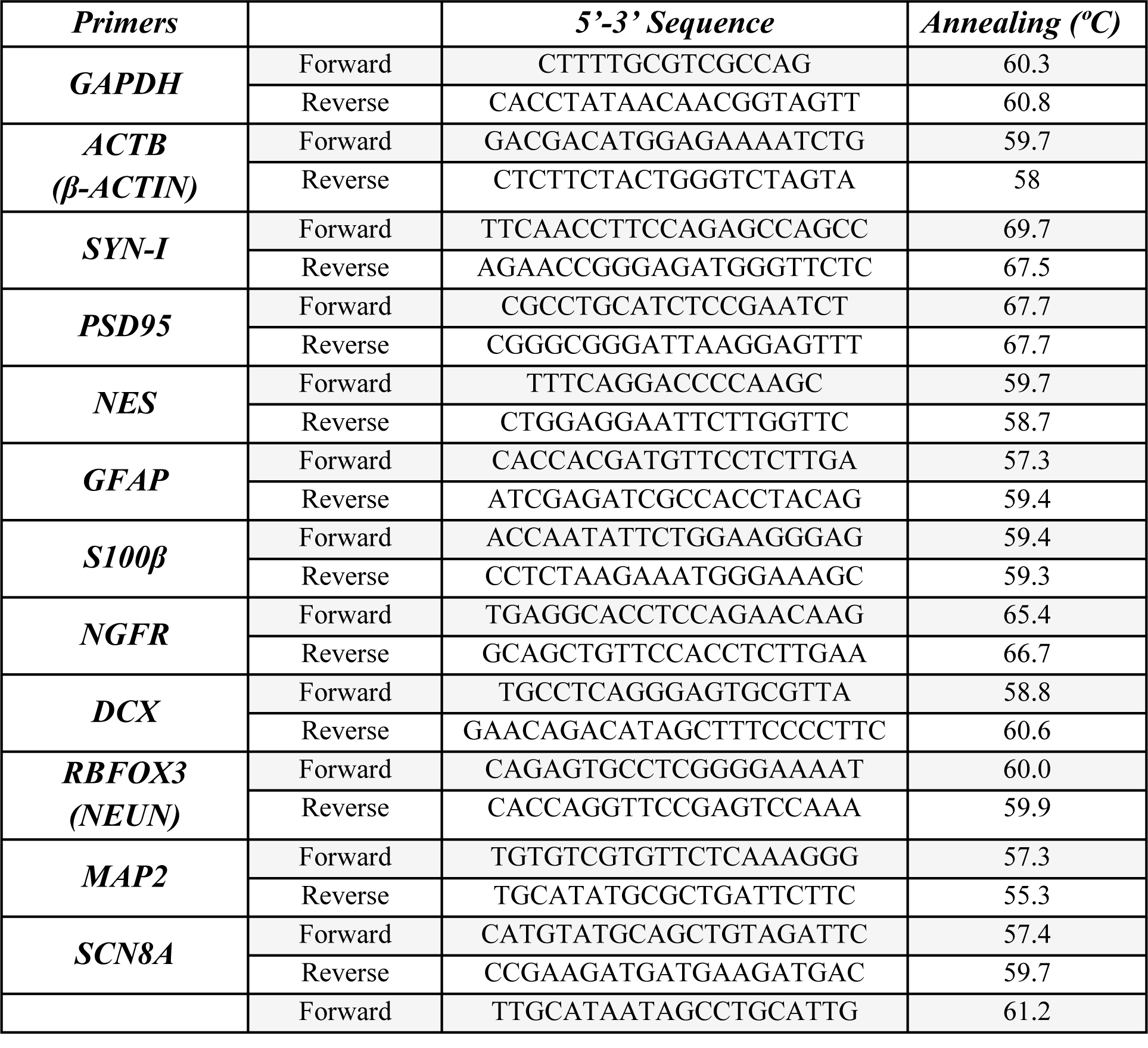
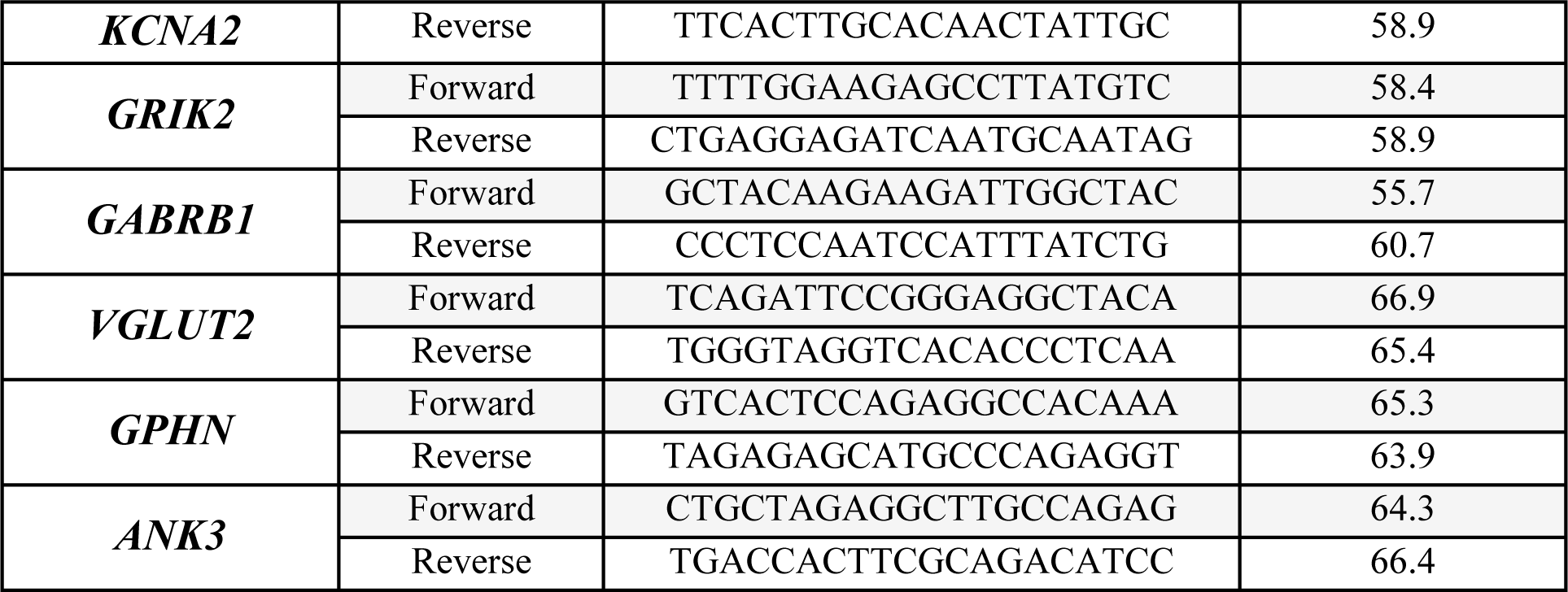
Primer pairs sequences used in this study.

### Electrophysiology

Patch-clamp experiments were performed in hDPSCs from different donors differentiated for 2-months. Whole-cell current-clamp and voltage-clamp measurements were performed at room temperature using MultiClamp 700B amplifier (Molecular Devices, San Jose, CA). Signals were filtered at 1 kHz and acquired at a 10 kHz sampling rate using a DigiData 1550 data acquisition system and pCLAMP 10.3 software (Molecular Devices, San Jose, CA). The bath solution contained (in mM): 140 NaCl, 5 KCl, 2 MgCl_2_, 10 HEPES-NaOH, 2 CaCl_2_ and 10 glucose (pH=7.3–7.4 adjusted with NaOH). Patch clamp pipettes were made of borosilicate glass (Sutter Instruments) and had a resistance of 4–6 MΩ with an internal solution containing (in mM): 135 potassium gluconate, 10 KCl, 10 HEPES, 1 MgCl_2_, and 2 ATP-Mg (pH=7.3adjusted with KOH). Fast and slow whole-cell capacitances were neutralized, and series resistance was compensated (∼70%).

Coverslips containing hDPSCs were placed on a chamber and cells were visualized using infrared-differential interference contrast optics (Olympus BX51WI microscope, Olympus Optical, Japan) and 40X water immersion lens. To measure the different currents, cells were held at −70 mV. Sodium current (I_Na_) was generated by 10-ms depolarizing pulses from −60 to +60 mV every 15 s with 10 mV steps and potassium currents (I_K_) by the application of 400 ms depolarizing pulses from −10 mV to +180 mV every 20 s with 10 mV steps. I_K_ and I_Na_ amplitudes were measured at the peak outward and inward values, respectively. We used a pump perfusion system (2 ml/min) to selectively block I_Na_ with 1 μM tetrodotoxin (TTX).

To measure spontaneous cell activity in voltage-clamp or current-clamp mode, cells were held at a −70 mV holding potential and recorded for 2 minutes. In current-clamp mode, action potentials (APs) were evoked by 3 consecutive current injections of 1500 ms duration, each separated by a 1000 ms interval. This protocol was repeated 6 times, with the current injection increasing by 30 pA with each iteration (see **Supplementary video 3**). Data from current-clamp and voltage-clamp experiments were analyzed using Clampfit software (Molecular Devices, CA, USA).

### Statistical analysis

For statistical analysis, SPSS 28.0 (Armonk, NY, USA) and GraphPad Prism 5 software (Boston, MA, USA) were used. According to the characteristics of the data distribution, comparisons between two groups were made via either Student’s t-test or the Mann‒Whitney U test. P values less than 0.05 were considered statistically significant. The results are presented as means ± SEM.

## Results

### FBS-induced changes influence the long-term neurodifferentiation fate of hDPSCs

*In vitro* culture media environment cues are decisive in the effectiveness of hDPSCs differentiation into neural cells. The ability of hDPSCs to differentiate into non mesenchymal lineages is compromised by the presence of fetal serum in the culture medium [28,29]. Furthermore, the creation of a neurosphere-like structure is critical for establishing close physical contacts between neural progenitors, which is necessary for neural fate commitment [30,31]. Thus, two distinct differentiation procedures were used to assess the reversibility of FBS-induced changes and the influence of the microenvironment generated inside the dentosphere on the hDPSC neurodifferentiation process. When hDPSCs were grown in serum-containing proliferation medium (**Fig. 1a**), they formed an adherent cell monolayer, with cells acquiring a characteristic mesenchymal spindle-like morphology even after being thereafter switched to a neural differentiation medium for 21 days (**Fig. 1c**). In contrast, cells initially cultured as floating spheres (**Fig. 1b**) in a serum-free proliferation medium —commonly used for brain NSC expansion— generated cells with a neural-like dendritic morphology, with smaller cell bodies and very long, thin and ramifying processes when they were switched to the same differentiation medium (**Fig. 1c**).

**Figure. 1:**
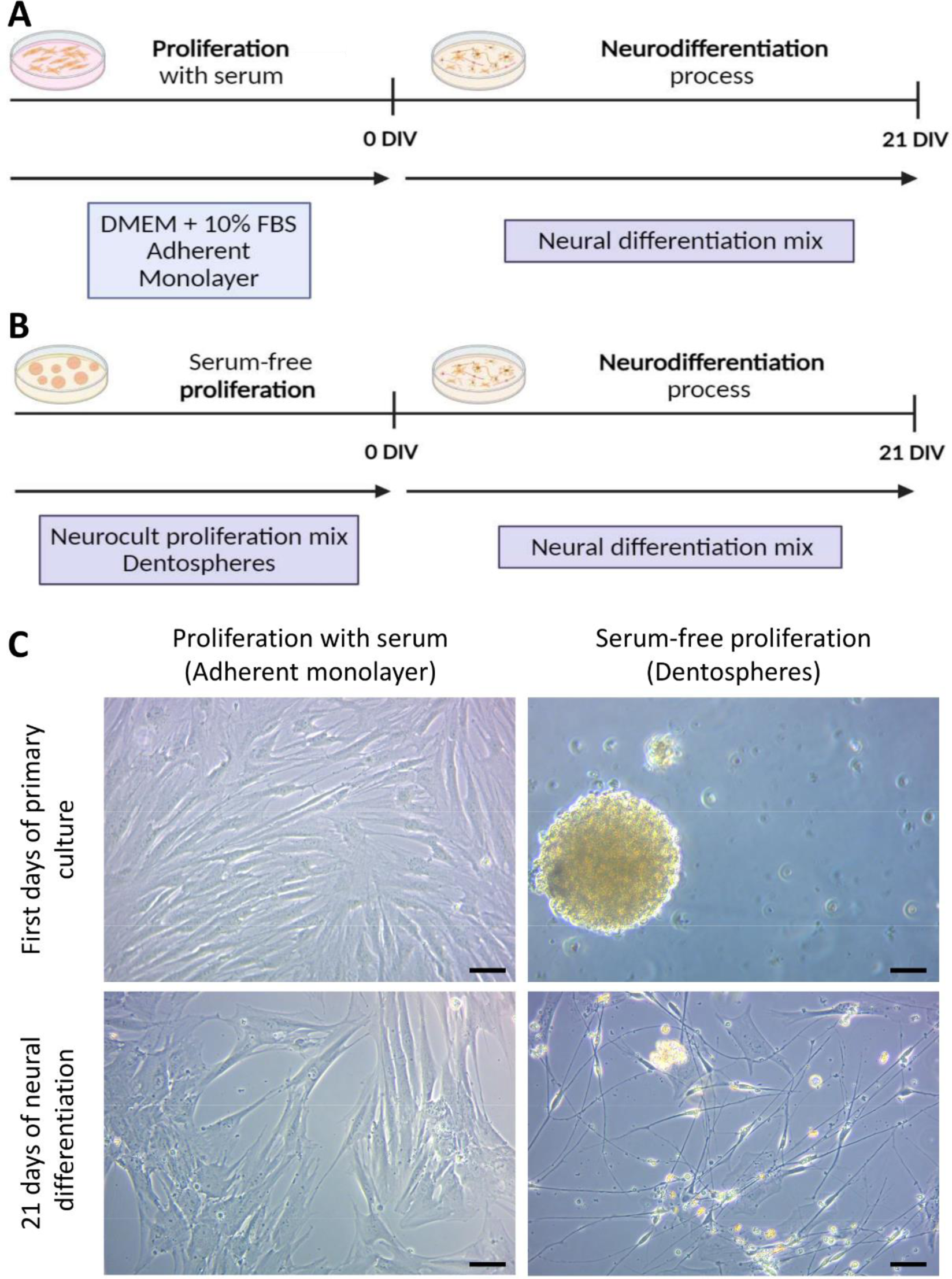
Differences between the conducted protocols for hDPSCs neurodifferentiation. **(A)** Scheme of hDPSCs that were cultured in DMEM with 10% FBS as an adherent monolayer and then switched for 21 days to a neural induction mix. **(B)** hDPSCs grown as floating dentospheres in serum-free neurocult proliferation mix and then switched for 21 days into a neural differentiation mix. **(C)** Bright field images of primary cultures performed with serum and without it and their appearance after switching them for 21 days to the same neural differentiation mix. Scale bar: 50 μm.

### The expression of neural stem cell markers decreases in hDPSCs after the neural induction process

Certain cytoskeletal proteins including the intermediate filaments Nestin and GFAP, which are frequently employed as NSC markers, are abundantly expressed in hDPSC cultures [12,13]. After three weeks of *in vitro* growth and prior to the neural induction process, we were able to detect both markers by immunofluorescence under each of our experimental conditions (**Fig. 2a-b**, **Fig. 3a-b**). Nevertheless, an increased number of Nestin (90.6±2.7% in serum-free vs 77.9±3.4% in serum; p=0.0007) and GFAP (96.7±1.0% in serum-free vs 87.0±3.5% in serum; p=0.0044) positive cells were detected in hDPSCs grown as floating spheres in comparison with those grown as monolayers in the presence of serum, which may suggest that the microenvironment within the spheres is more suitable for the development and maintenance of neurogenic cells *in vitro*.

**Figure 2:**
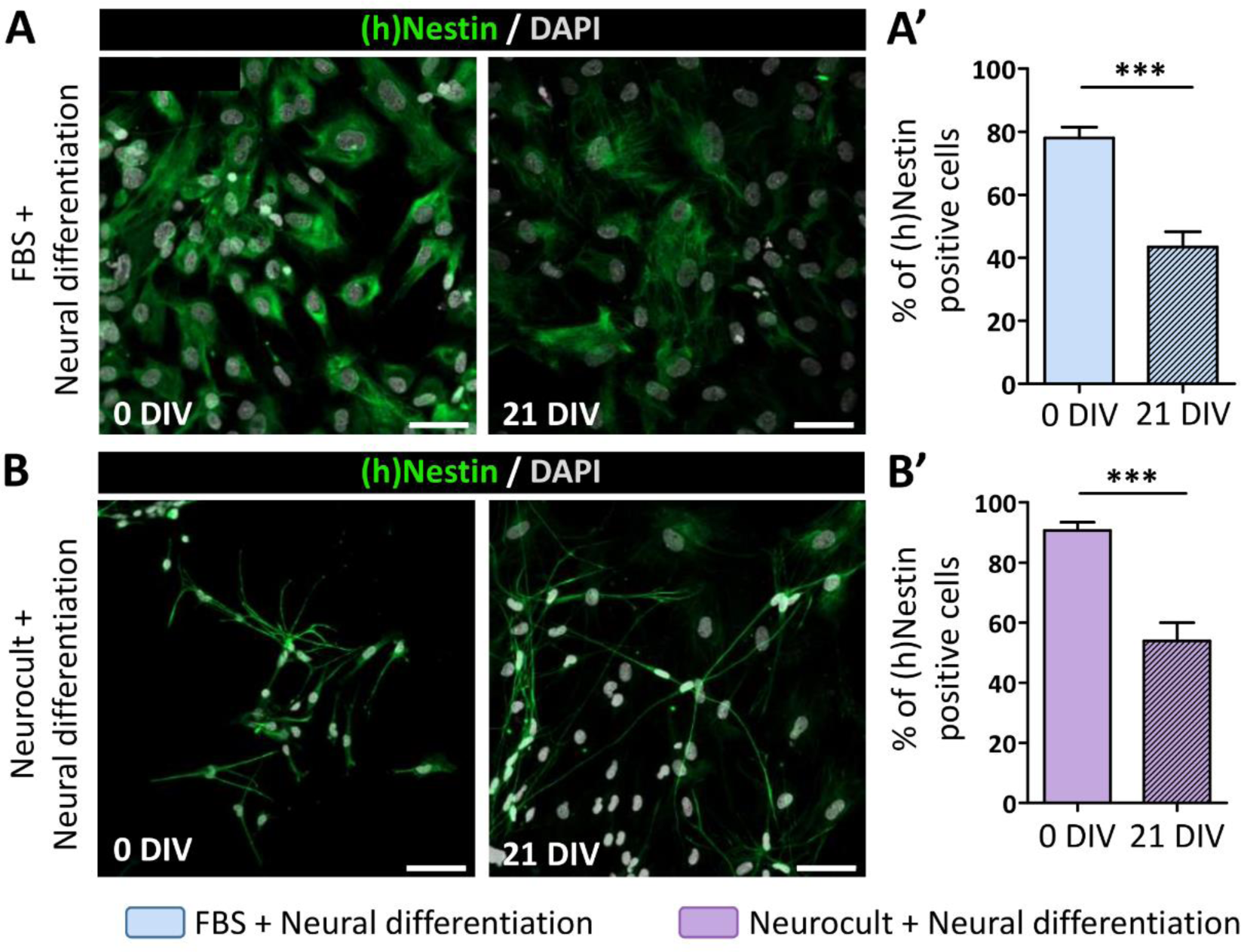
Human Nestin ((h)Nestin) stem cell marker expression decrease after 21 days of neurodifferentiation. **(A)** hDPSCs proliferated in serum containing media, showing fibroblast like morphologies and a detectable decrease of (h)Nestin labelling (green) after 21 days of differentiation. Scale bar 50 μm. **(A’)** Quantification of (h)Nestin (green) positive cells just after switching them from a DMEM-10% FBS media to a neural induction mix and 21 days afterwards. ***p<0.001 **(B)** Cells grown in neurocult proliferation mix presenting a ramified morphology and a decrease in (h)Nestin positive labelled cells (green) after 21 days of differentiation. All images are counterstained with DAPI. Scale bar 50 μm. **(B’)** Percentage of (h)Nestin positive cells after switching them from a serum-free proliferation mix into a neural induction mix and 21 days afterwards. ***p<0.001. U-Mann Whitney (Two-tailed) test.

**Figure 3:**
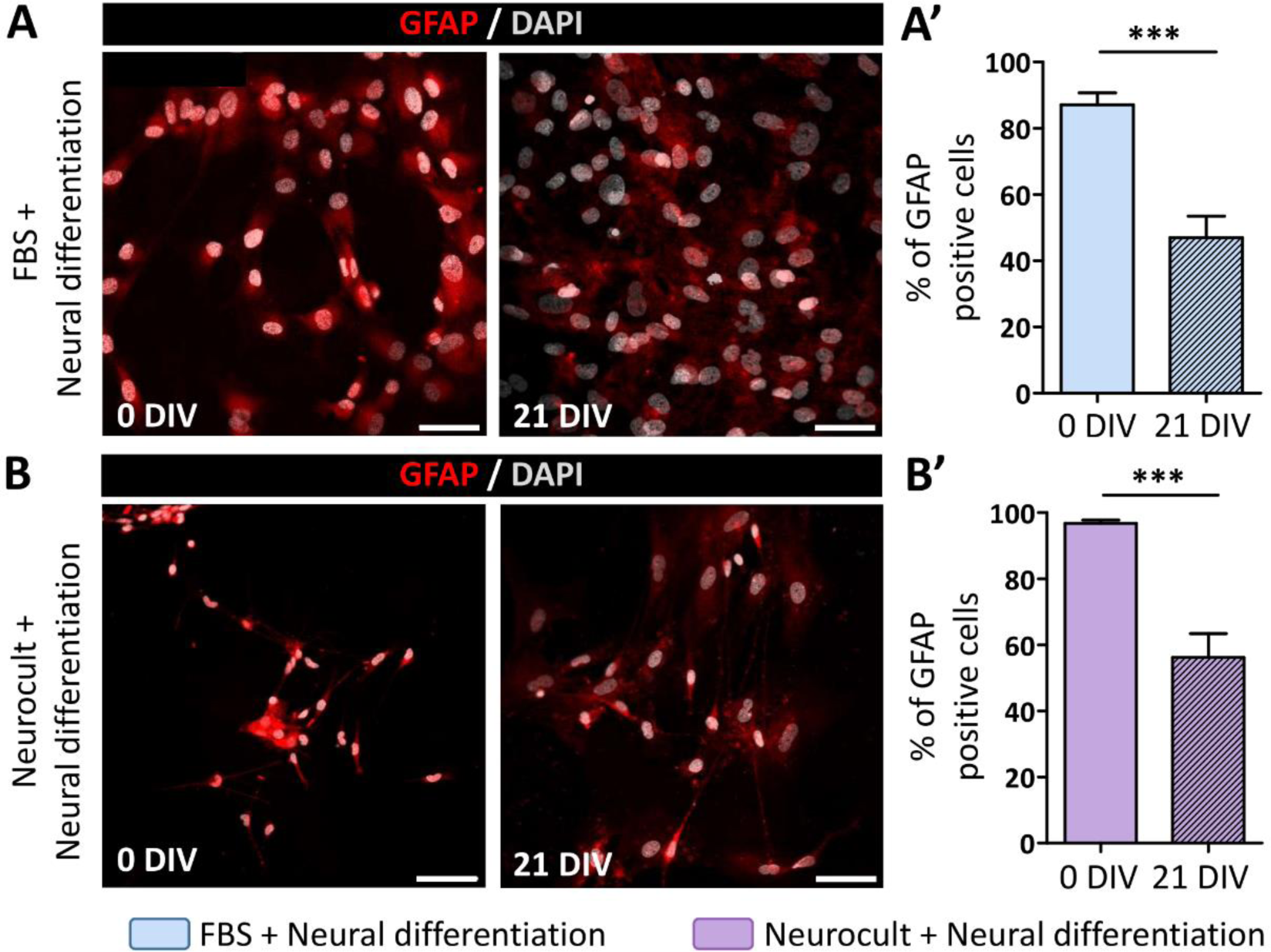
Glial fibrillary acidic protein (GFAP) expression decrease in hDPSCs after 21 days of neural induction. **(A)** hDPSCs grown with serum stained with GFAP antibody (red) at the beginning of the neural differentiation process and 21 days thereafter. Scale bar 50 μm. **(A’)** Graph showing the decrease of GFAP positive cells in hDPSCs expanded with DMEM-10% FBS after switching them to a neural induction mix. ***p<0.001 **(B)** Fluorescence photomicrographs of GFAP (red) in hDPSCs expanded in serum-free Neurocult proliferation mix just after changing them to a neural induction mix and 21 days after. All images are counterstained with DAPI. Scale bar 50 μm. **(B’)** Graph showing GFAP positive cell percentage reduction 21 days after switching cultures from Neurocult proliferation mix to neural differentiation media. ***p<0.001. U-Mann Whitney (Two-tailed) test.

During the natural neurodifferentiation process, these immature markers undergo downregulation to progressively be replaced by adult neuronal markers [32]. To test whether our neuroinduction protocols affected the expression of immature NSC markers, we assessed the expression of Nestin and GFAP at the beginning of the differentiation process and 21 days afterwards (21DIV). At 21 days of differentiation, the percentage of immunopositive cells was similarly reduced for both markers in both neurodifferentiation protocols thus suggesting that stemness of hDPSCs is equivalently reduced both in the presence (**Figs. 2a-a’, Fig. 3a-a’**) or absence (**Figs. 2b-b’, Fig. 3b-b’**) of serum. However, it is noteworthy that cells grown with FBS as an adherent monolayer still presented mesenchymal-like morphologies even after they were switched to neural induction media (**Fig. 2a**, **Fig. 3a**).

### Neural differentiated hDPSC cultures generated mature neuronal and glial cells

Considering the decrease in hDPSCs stemness after neural induction protocols, we characterized the marker expression pattern of the differentiated cell population. S100β, a mature glial marker [33] whose expression in GFAP-expressing cells coincides with their loss of stemness and the maturation of the astroglial phenotype [34], is also a characteristic marker of certain peripheral nervous system cells. Specifically, its co-expression with the low-affinity neurotrophin co-receptor p75^NTR^ constitutes a characteristic Schwann cell molecular profile [35]. After 21DIV, the expression of S100β and p75^NTR^ was detected by immunocytochemistry under each of our culture conditions suggesting that a fraction of the differentiated cells had committed to peripheral glia (**Fig 4d**). As shown in the Venn diagrams, 2.5% of the cells that were immunoreactive for S100β also colocalized with p75^NTR^, indicating a Schwann cell phenotype, in hDPSCs grown with FBS **(Fig. 4a)**. However, this phenotype was preferentially acquired by those cells that had initially been grown in the absence of serum, in which we detected a 9% of S100β^+^/p75^+^ cells (**Fig. 4b**). Overall, hDPSCs grown in the presence of serum presented a significantly reduced percentage of S100β (4.1±1.7%) and p75^NTR^ (3.4±1.4%) positive cells compared with those grown in serum-free dentosphere conditions, whose percentages were 10.8±3.1% and 14.1±3.5% respectively (S100β: p < 0.0077 and p75^NTR^: p = 0.0011, two-tailed Mann‒Whitney U test; **Fig. 4c**).

**Figure 4:**
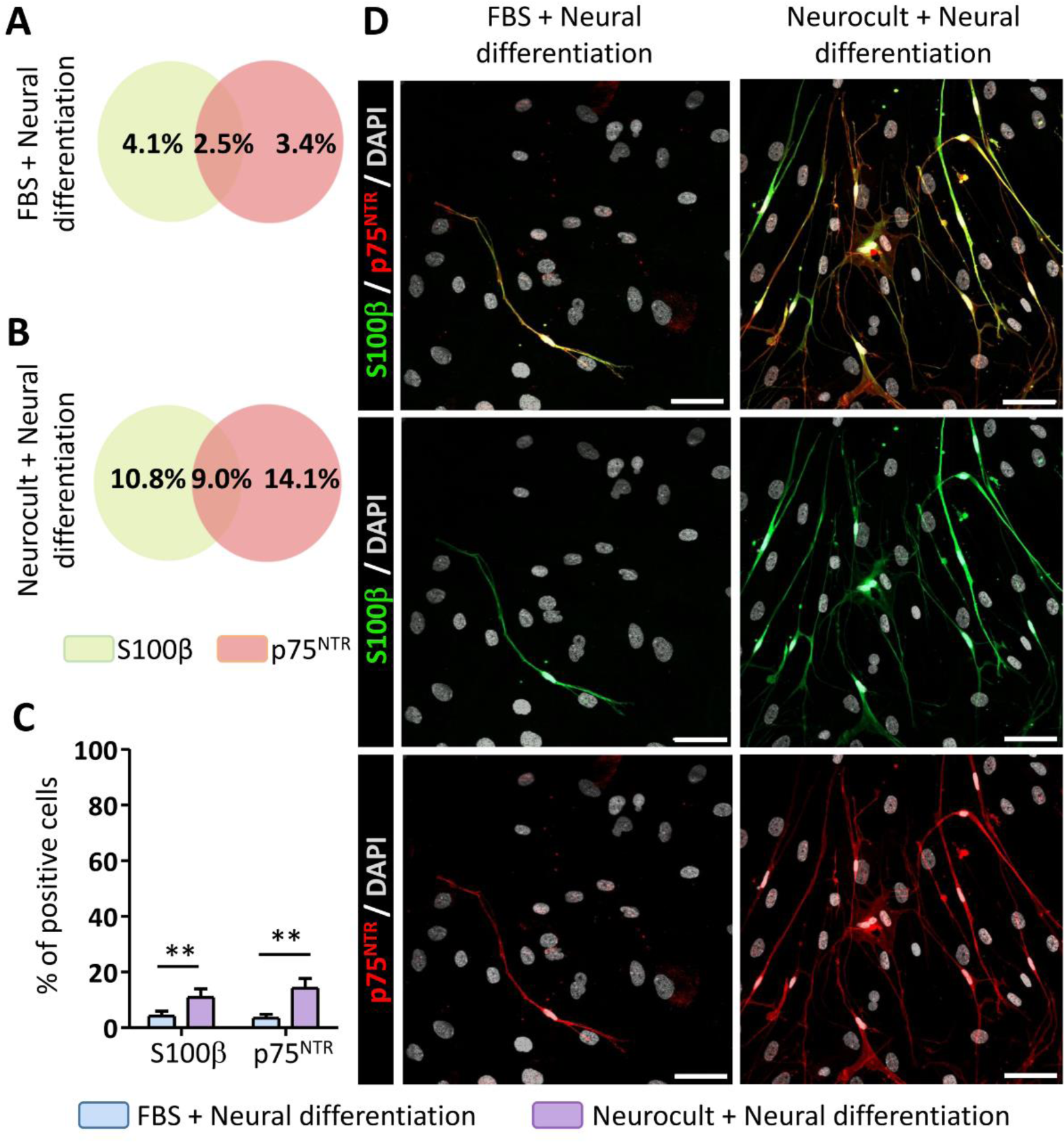
Neurodifferentiated hDPSCs are able to commit toward Schwann Cell phenotypes. Venn’s diagrams representing the percentage of positive cells for S100β and p75 glial markers after 21 days differentiation, as well as the percentage of co-localizing cells for hDPSCs grown with FBS **(A)** and those that were not **(B). (C)** Graphs showing the percentage difference for S100β and p75 markers between hDPSCs that have grown with FBS and those in serum-free Neurocult media after switching them to the same neural induction mix during 21 days. **p<0.01 **(D)** Higher rates of immunopositive cells for p75 (red) and S100β (green) could be observed in hDPSCs grown with Neurocult comparing to those that were grown in FBS after 21 days of differentiation. Scale bar 50 μm. Statistical analyses were conducted by U-Mann Whitney (Two-tailed) test.

The presence of a neuronal-like molecular phenotype was assessed by the immature neuronal marker doublecortin (DCX) and the mature neuronal nuclear protein (NeuN). Both markers were shown to be expressed by hDPSCs under both experimental conditions (**Fig. 5a**).

**Figure 5:**
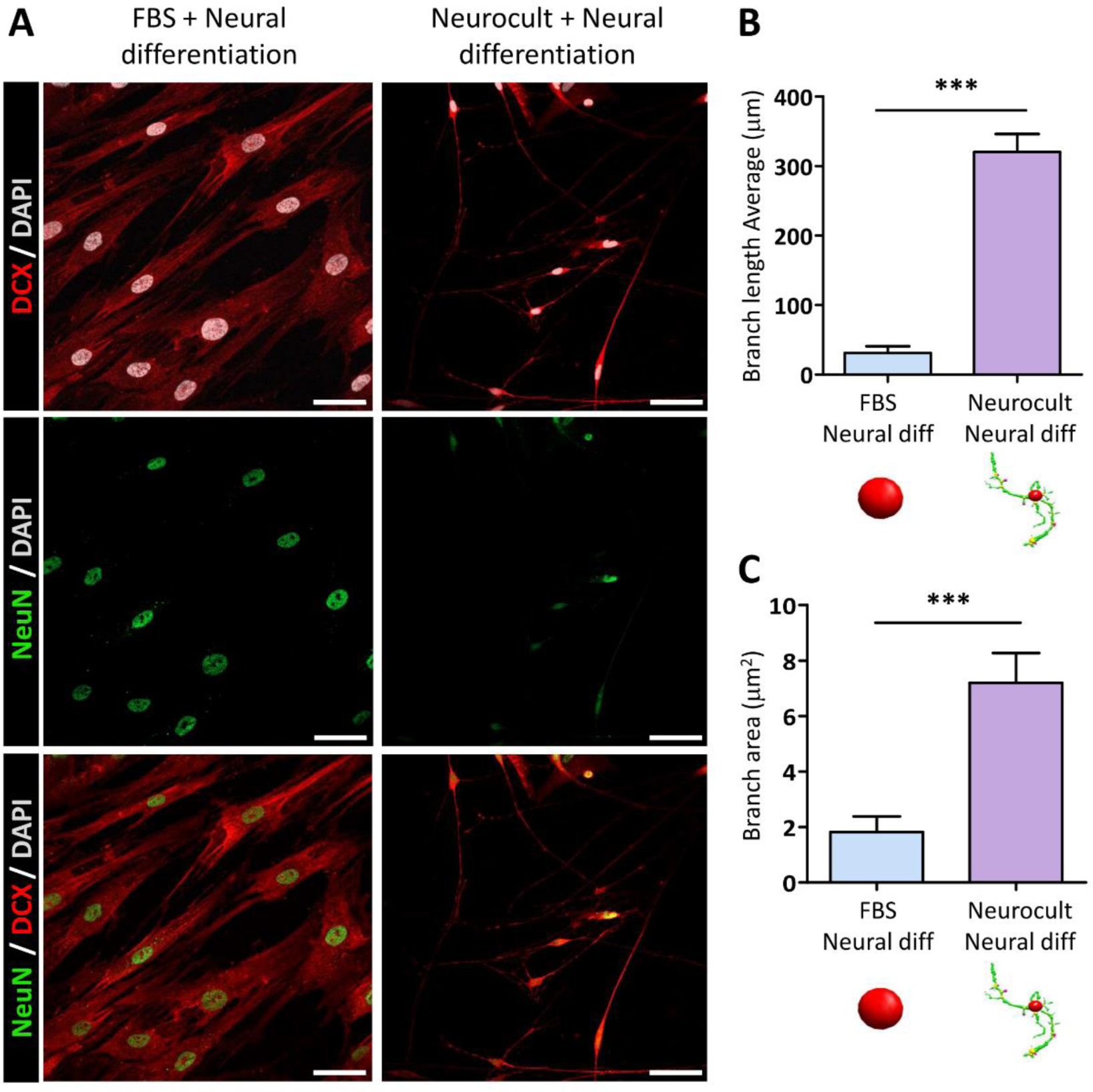
Neuronal marker expression and cell morphological analysis after 21 days of neurodifferentiation. **(A)** Immunofluorescence images showing doublecortin (DCX) and neuronal nuclear protein (NeuN) positive labelling in hDPSCs and different morphologies observed between cells grown in a serum containing medium and those that were grown in a serum-free medium. Scale bar 50 μm. 3D Sholl-analysis of 30 cells per condition revealed longer branch lengths measured in μm **(B)** and a larger overall surface (μm2) **(C)** occupied by hDPSCs processes in those cells previously grown as a floating dentospheres. ***p<0.001. Statistical analysis conducted by U-Mann Whitney (Two-tailed) test.

Nevertheless, the nuclear shape and cell morphology were notably different between the conditions. Indeed, hDPSCs previously cultured with FBS exhibited prominent spindle-like cytoplasmic morphologies resembling those of fibroblasts and an increased nuclear area (**Supplementary Figure 1**).

**Supplementary Figure 1:**
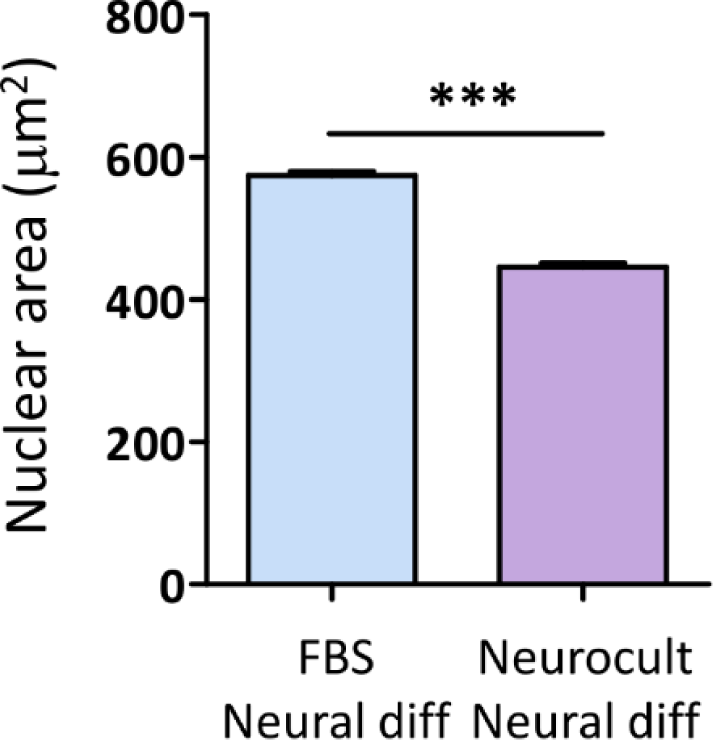
Nuclear area measured after 21 days of neural induction process. Graph showing a bigger nuclear area (μm2) in cells cultured with FBS after measuring 1200 nuclei per experimental condition with the NII plugin for Image J. ***p<0.001. U-Mann Whitney (two-tailed) test.

On the other hand, hDPSCs initially cultured as spheres presented reduced nuclear areas, smaller cell bodies and many thin ramifications. These morphological differences were characterized via 3D-Sholl analysis, and we detected significantly longer branches (320.2±26.0 μm) (**Fig. 5b**) covering a larger surface (7.2±1.0 μm^2^) (**Fig. 5c**) in hDPSCs cultured in the absence of serum (p < 0.0001, two-tailed Mann‒Whitney U test). Despite the positive DCX and NeuN immunolabeling (**Fig. 5a**) and the observed neuronal-like morphologies in those cells that previously expanded as dentospheres (**Fig. 5b-c**), we could not obtain any electrophysiological recordings showing that they were electrically excitable cells (**Table 2**).

**Table 2:** Electrophysiological recordings.

| DIV | Neurocult-Neural diff |  |  |  |  |  | Neurocult-Neural diff KCl |  |  |  |  |  |
| --- | --- | --- | --- | --- | --- | --- | --- | --- | --- | --- | --- | --- |
|  | I<sub>Na</sub> |  | I<sub>K</sub> |  | AP |  | I<sub>Na</sub> |  | I<sub>K</sub> |  | AP |  |
|  | Trial | Resp. | Trial | Resp. | Trial | Resp. | Trial | Resp. | Trial | Resp. | Trial | Resp. |
| 21 | 5 | 0 | 3 | 1 | 3 | 0 | 20 | 3 | 13 | 3 | 15 | 0 |
| 60 |  |  |  |  |  |  | 24 | 5 | 21 | 3 | 29 | 6 |

### hDPSCs expressed mature neuronal and synaptic markers after the neural differentiation protocol was improved with RA and KCl

As reported in the literature, RA is known to improve NSC survival and favor differentiation into neuronal phenotypes, whereas repeated KCl depolarizations reduce cell proliferation together with an increase in neurite outgrowth [23]. Thus, we went one step ahead in improving the neural differentiation protocols by adding 10 μM RA and one hour depolarizing pulses of 40 mM KCl every two days to the neural induction mixture (**Fig. 6a**).

**Figure 6:**
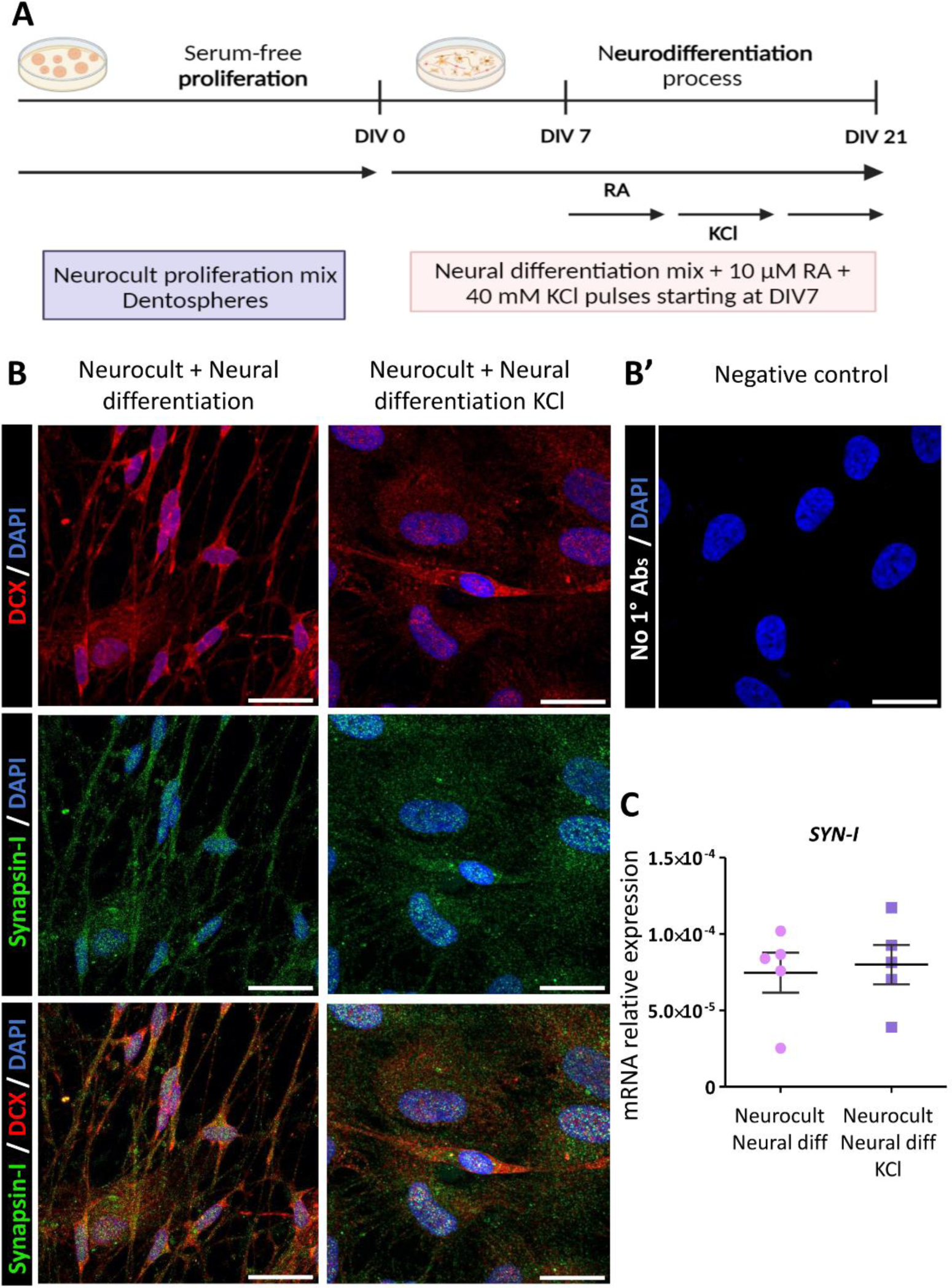
hDPSCs differentiation process strengthening with retinoic acid (RA) and KCl and Synapsin-I pre-synaptic marker expression. **(A)** Neural induction process of those cells proliferated as a dentosphere was upgraded by adding 10 μM retinoic acid (RA) and one hour 40 mM pulses of KCl into differentiation media, starting at day seventh of differentiation. **(B)** Confocal images showing positive labeling for the presynaptic protein Synapsin-I (green dots) in those cells treated with RA and KCl and the untreated ones. Scale bar 20 μm. **(B’)** Synapsin-I expression at mRNA level measured in cultures from hDPSCs obtained from 5 different donors in each experimental condition.

At 21DIV, we assessed the transcript expression levels of different stem (*NES*), glial (*GFAP*, *S100B*, *NGFR*) and neuronal (*DCX*, *NEUN*, *MAP2*) marker genes and we did not observe significant differences between cells differentiated in the presence or absence of RA and KCl (**Supplementary Figure 2**).

**Supplementary Figure 2:**
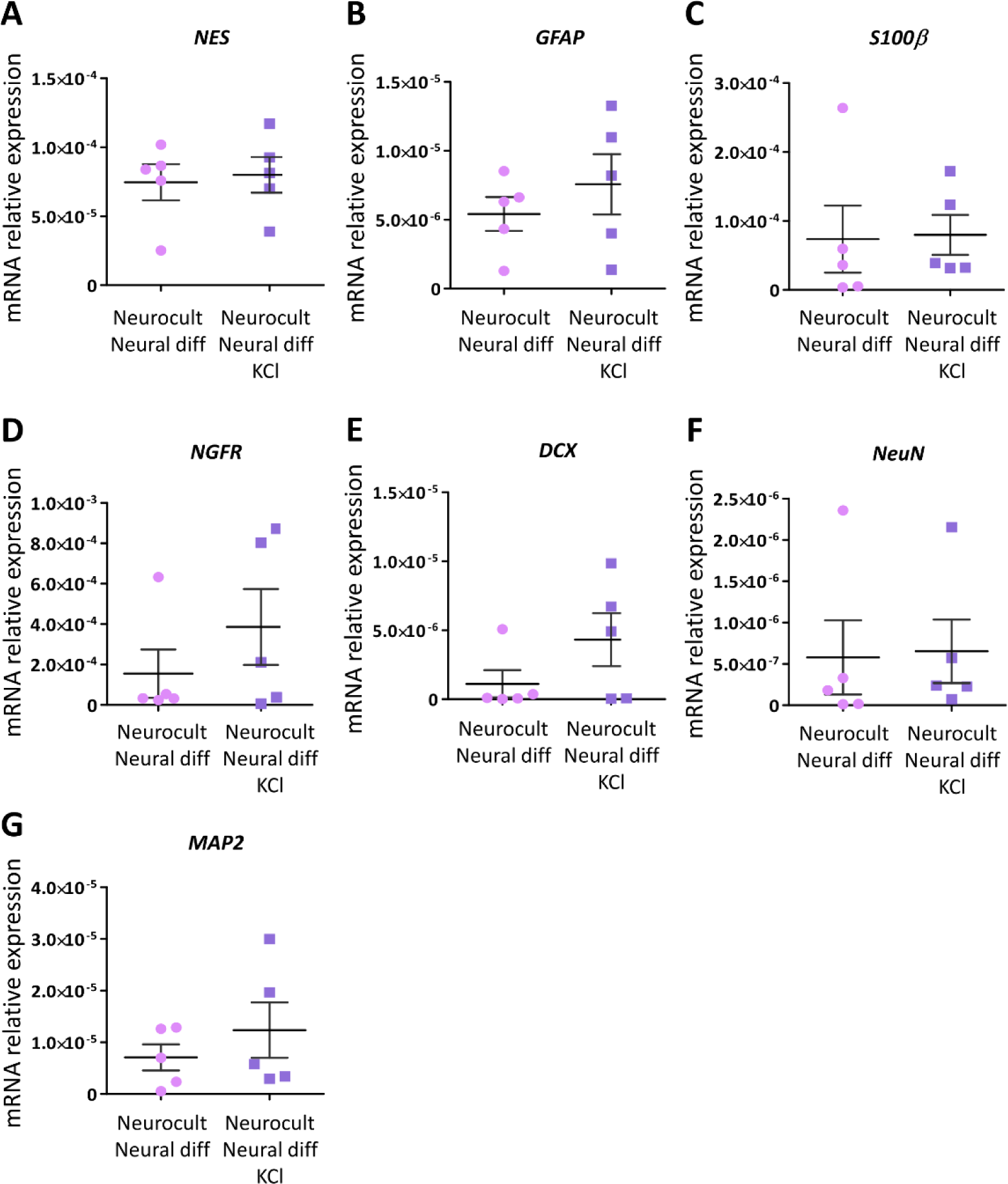
mRNA expression levels after RA and KCl addition to Neurocult differentiation mix at 21 days of neurodifferentiation. RT‒qPCR analysis of **(A)** Nestin, **(B)** GFAP, **(C)** S100β, **(D)** NGFR and neuronal **(E)** DCX, **(F)** NeuN and **(G)** MAP2 relative mRNA expression in hDPSCs treated or not with RA and KCl.

Moreover, the expression of Synapsin-I, a protein present in synaptic vesicles and implicated in synaptogenesis [36], was also detected both in those cultures treated with KCl and RA and in those that were not (**Fig. 6b-6c**).

### hDPSCs expressed components of both excitatory glutamatergic synapses and inhibitory GABAergic synapses after the neural induction process

To further characterize the heterogeneity of the resulting cell population obtained after the neurodifferentiation process, we assessed the expression of different proteins required to construct excitatory glutamatergic synapses, as well as inhibitory GABAergic synapses. Differentiated hDPSCs presented a positive punctate immunolabeling signal corresponding to the postsynaptic density protein PSD95 (**Fig. 7a**), which is a central organizer of postsynaptic complexes in glutamatergic synapses [37]. These results were confirmed by qRT‒ PCR (**Fig. 7a’**). On the other hand, the presence of a glutamate vesicular transporter (VGlut2) was also detected **(Fig. 7b**). These results could indicate the capacity of some of the differentiated cells to package glutamate into synaptic vesicles. Nevertheless, the ionotropic glutamate receptor subunit kainate type 2 (*GRIK2*) was significantly overexpressed in hDPSCs treated with KCl and RA compared with non treated cultures (p = 0.016, two-tailed Mann‒Whitney U test; **Fig. 7c**). With regard to inhibitory synapses, sequential exposure to RA and KCl has been previously demonstrated to commit NSCs toward a GABAergic phenotype after neural differentiation [23]. Interestingly, after 21DIV, we detected GABA-positive cells under our culture conditions, with a markedly higher intensity of labeling in those cells where the differentiation was strengthened with KCl and RA (**Fig. 8a-a’-a’’**). Differentiated hDPSCs positive for GABA also expressed glutamic acid decarboxylase GAD65, a protein responsible for GABA synthesis, which may indicate that differentiated hDPSCs possess the molecular machinery for the production of this inhibitory neurotransmitter (**Fig. 8a-a’**).

**Figure 7:**
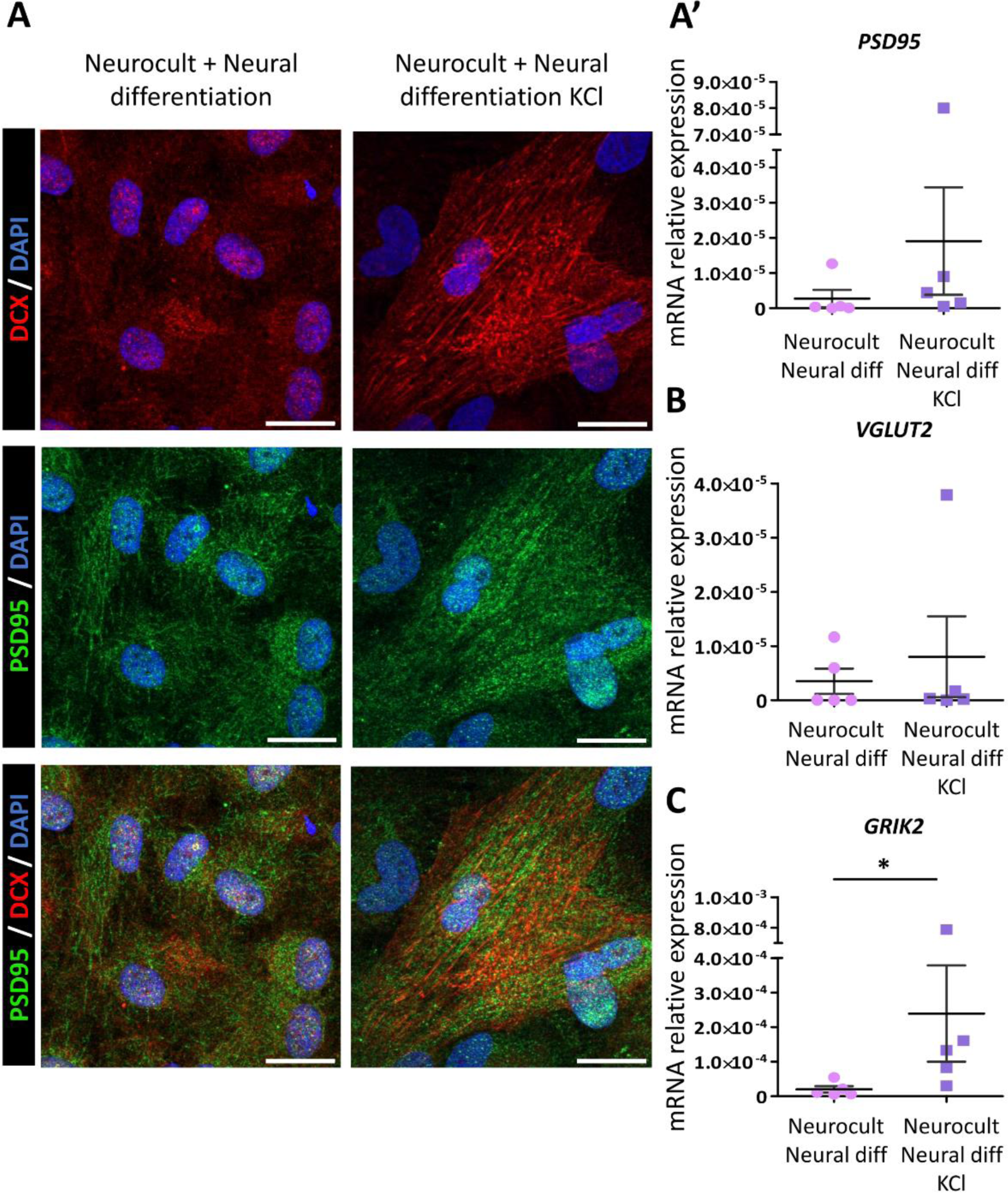
Excitatory glutamatergic synapse components in hDPSCs after 21 days with or without RA and KCl. **(A)** Inmunofluorescence images showing a positive signal for the postsynaptic PSD95 protein (green dots) either in hDPSCs whose neural differentiation was strengthened with RA and KCl and those that not. Scale bar 20 μm. **(A’)** PSD95 and vesicular glutamate transporter (vGLUT2) **(B)** expression measured at mRNA level in cultures performed for each experimental condition. **(C)** Graph showing an overexpression of the glutamate ionotropic receptor kainate type subunit 2 (GRIK2) in hDPSCs treated with RA and KCl *p<0.016. Statistical analysis performed by U-Mann Whitney (Two-tailed) test.

**Figure 8:**
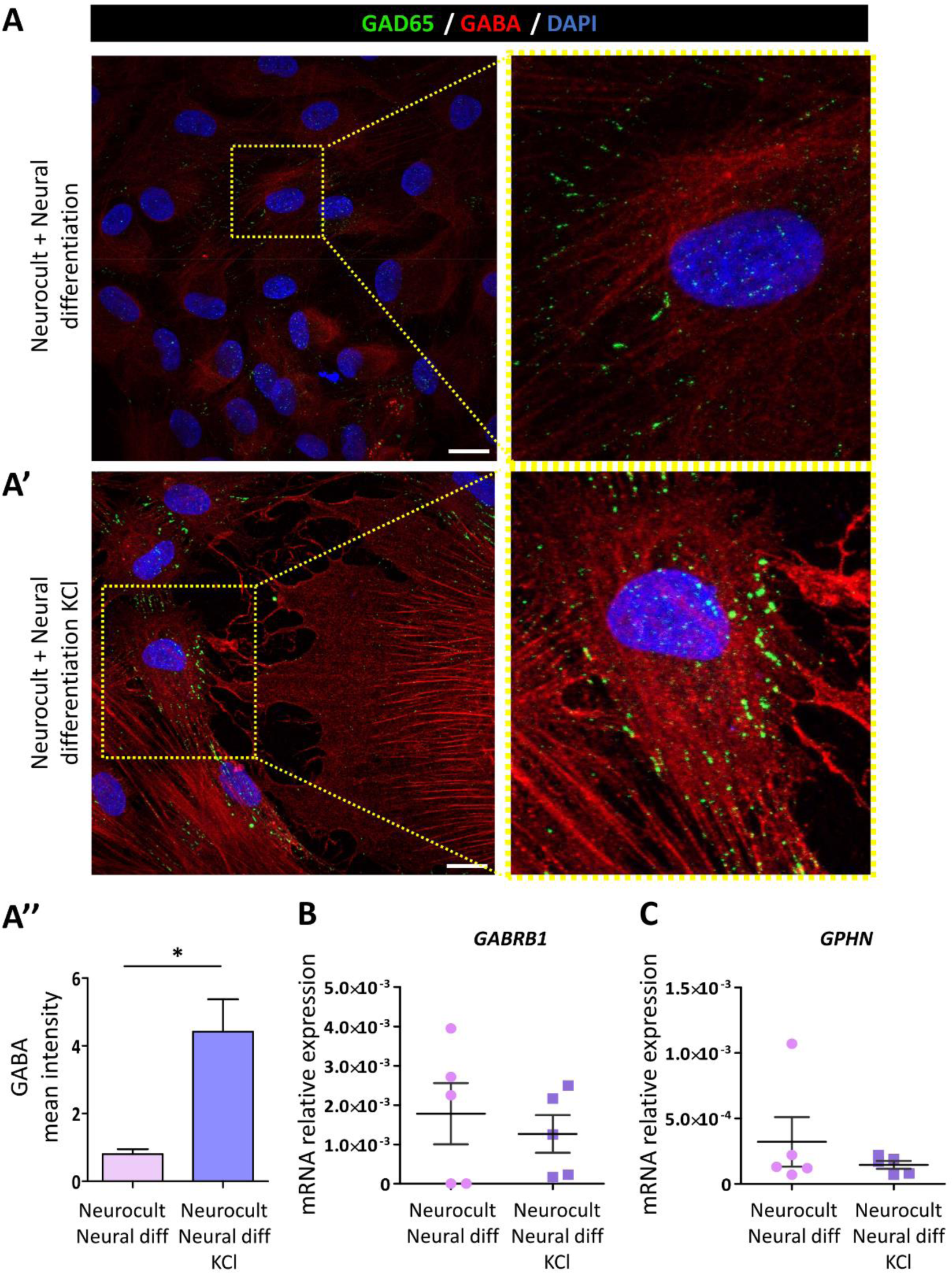
Inhibitory GABAergic synaptic proteins in hDPSCs after 21-day differentiation with or without RA/KCl treatment. GABA neurotransmitter and glutamate descarboxylase (GAD-65) (green dots) inmuno positive labeling in cells neurodifferentiated without RA and KCl **(A)** and more intense labeling of those GABAergic markers in those cells that were treated **(A’).** Scale bar 20 μm. **(A’’)** Graph showing higher GABA labeling intensity in hDPSCs treated with KCl and RA * p<0.05. Gamma-aminobutyric acid type A receptor subunits (GABRB1) **(B)** and gephyrin (GPHN) **(C)** mRNA relative expression in hDPSCs differentiated with the same neural differentiation mix and those in with RA and KCl. U-Mann Whitney test (Two-tailed).

Furthermore, the mRNAs encoding the gamma-aminobutyric acid type A receptor subunit (*GABRB1*) (**Fig. 8b**) and the central GABAergic synapse organizer Gephyrin (*GPHN*) (**Fig. 8c**) were also detected via qRT‒ PCR in our differentiated hDPSCs.

### hDPSCs grown in the presence of RA and KCl display an electrophysiologically excitable neuron-like phenotype

In mature neurons, the distal end of the axon initial segment (AIS) is a specialized region, containing a high density of voltage-gated sodium channels (Na_v_) [38], where action potentials are shaped and initialized [39]. Ankyrin-G is a key membrane cytoskeletal linker for the structural organization of the AIS, as it is known to anchor Na_v_ channels into this compartment [40]. After 21DIV, hDPSCs grown with the upgraded KCl and RA protocol showed a positive immunolabeling signal against Ankyrin-G (**Fig. 9a**). In addition, *ANK3* expression was confirmed at the mRNA level via qRT‒PCR (**Fig. 9a’**). To characterize whether differentiated hDPSCs also expressed the machinery required for action potential generation and propagation, voltage-gated sodium (*SCN8A*) and potassium (*KCNA2*) channel subunits expressions were measured via qRT‒PCR. *SCN8A* and *KCNA2* were significantly overexpressed in those cells treated with KCl and RA, suggesting a greater density of these voltage-channels in treated hDPSCs (*SCN8A:* p = 0.025, two-tailed Student’s t-test; **Fig. 9b –** *KCNA2:* p = 0.008, two-tailed Mann‒Whitney U test; **Fig. 9c**).

**Figure 9:**
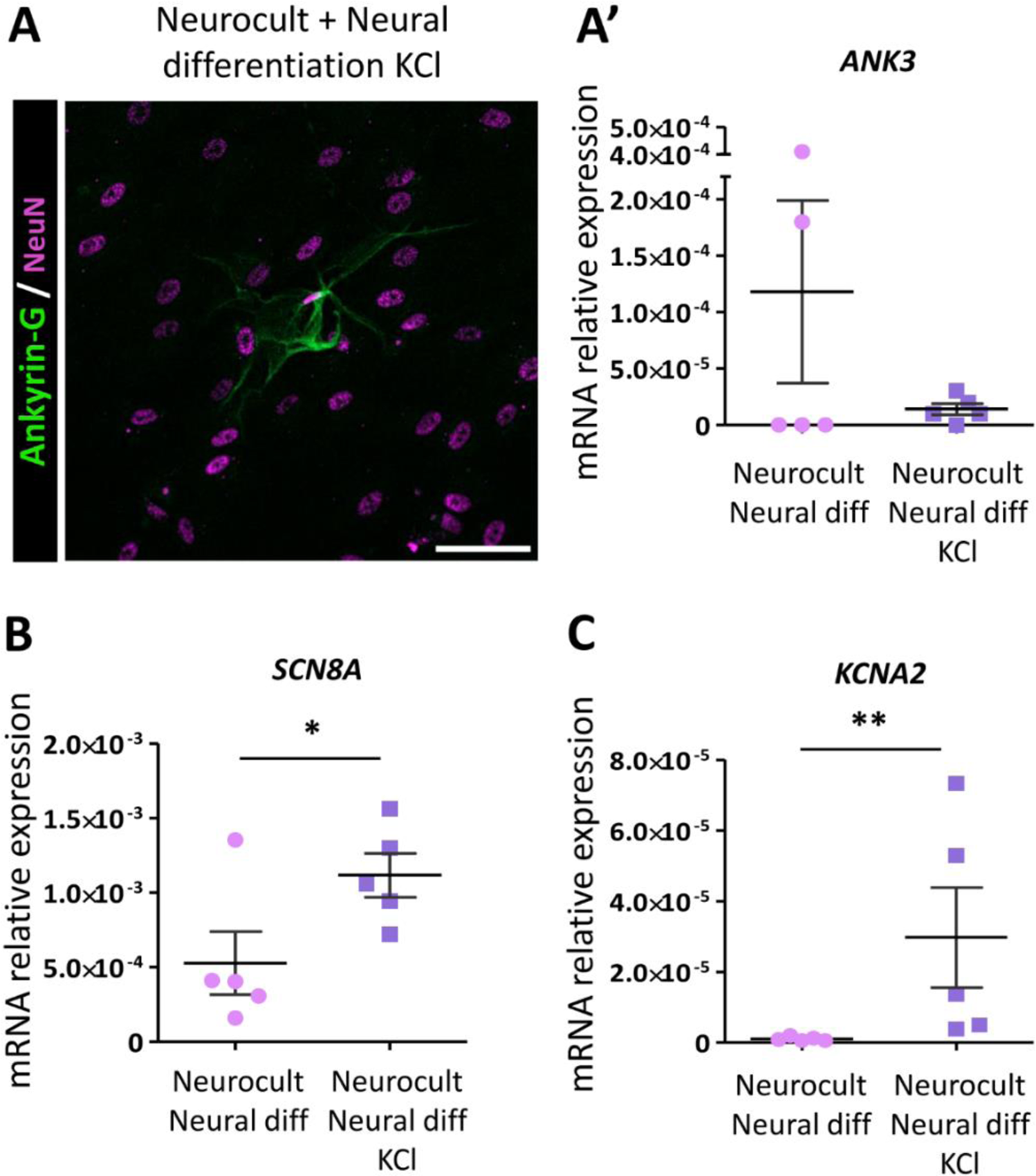
Ankyrin-G, SCN8A, and KCNA2 expression after 21 days in RA/KCl-treated and untreated hDPSCs. Axon initial segment Ankyrin-G protein (in green) and NeuN (in purple) inmuno positive labeling in hDPSCs differentiated with KCl and RA **(A).** Scale bar 50 μm. Ankyrin-G (ANK3) mRNA relative expression in hDPSCs differentiated with a standard neural differentiation mix and an upgraded mix with RA and KCl **(A’).** Graphs showing an increased expression of voltage-gated sodium (SCN8A) **(B)** and potassium (KCNA2) **(C)** channels in those cells treated with KCl and RA. * p<0.05, ** p<0.01. U-Mann Whitney test or Student’s t-test (Two-tailed).

The functionality of the KCl-RA treated and untreated hDPSCs was characterized by electrophysiology, but they did not generate action potentials despite their ability to generate voltage-dependent Na^+^ and K^+^ currents (**Table 2**). Consequently, we decided to take another step ahead and extended the hDPSCs differentiation process to two months (60DIV) to increase the frequency of mature neuronal-like cells. After neural induction with KCl and RA, we measured voltage-gated Na^+^ and K^+^ currents by electrophysiology. In neural-differentiated hDPSCs, sodium inward currents exhibited an activation threshold at approximately −30 mV, peaked at 0 mV (133 ± 41.4 pA) and exhibited a reversal potential at +60 mV (**Fig. 10a**). Furthermore, Na^+^-inward currents were completely blocked after perfusion of 1 μM tetrodotoxin (TTX) into the bath solution, confirming their generation by neuronal Na_v_ channels (**Fig. 10a**). Similarly, voltage-dependent potassium currents characterized by delayed-rectifier current-voltage (I-V) profile were also recorded. These currents were activated at membrane potentials above −10 mV and reached a peak value of 1339 ± 341 pA at 180 mV (**Fig. 10b**). Finally, single action potentials were elicited by depolarizing hDPSCs using a series of current injections. The initial protocol consisted of six current injections with increments of 30 pA and pulse durations of 2000 ms (**Supplementary video 1**). From 150 pA onwards, a single action potential was observed but additional increases in the current injection did not result in additional action potentials. To explore this further, we extended the pulse duration to 8000 ms, yet only one action potential was generated when the membrane potential exceeded −30 mV, the activation threshold for Na^+^ channels (**Supplementary video 2**). In another protocol, three depolarizing pulses of 1500 ms duration were applied, separated by 1000 ms intervals, and repeated six times with current increments of 30 pA. During each depolarization, a single action potential was generated, but the membrane had to return to its resting potential before another action potential could be initiated (**Fig. 10d & Supplementary video 3)**. Moreover, whole-cell voltage and current clamp recordings revealed that neurodifferentiated hDPSCs presented spontaneous activity in the absence of electrical stimulation (**Fig.10c**). Nevertheless, these cells did not show repetitive action potential firing patterns characteristic of fully mature neuronal cells after single current injections.

***Supplementary Video 1: APs of differentiated hDPSCs (Whole cell Current clamp):** Current clamp recordings after six current injections with increments of 30 pA and pulse durations of 2000 ms in two months differentiated hDPSCs treated with KCl and RA.*

***Supplementary Video 2: APs of differentiated hDPSCs (Whole cell Current clamp):** Current clamp recordings after six current injections with increments of 30 pA and pulse durations of 8000 ms in two months differentiated hDPSCs treated with KCl and RA.*

***Supplementary Video 3: APs of differentiated hDPSCs (Whole cell Current clamp):** Current clamp recordings after three depolarizing pulses of 1500 ms duration, separated by 1000 ms intervals, and repeated six times with current increments of 30 pA in two months neurodifferentiated hDPSCs treated with KCl and RA.*

**Figure 10:**
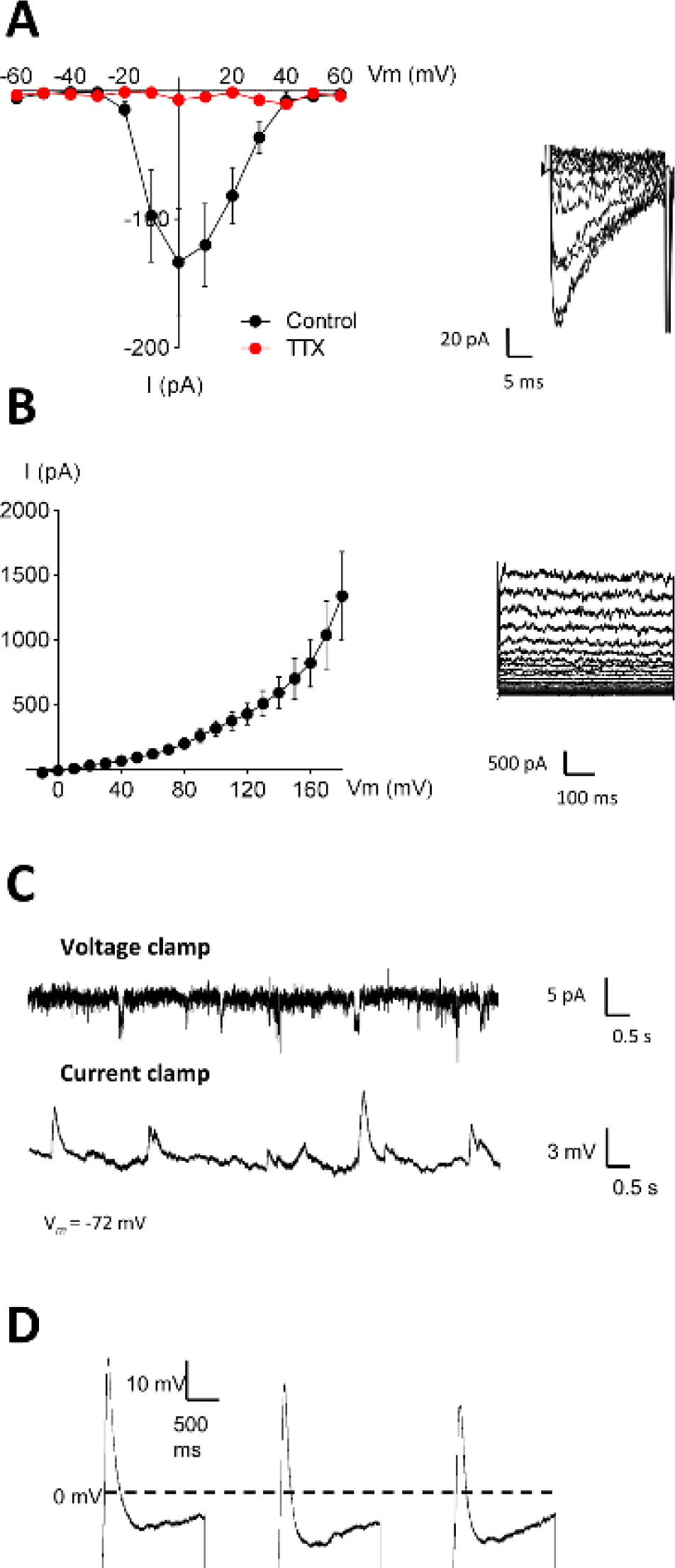
Electrophysiology of hDPSCs differentiated with KCl and RA. Neurodifferentiated hDPSCs displayed voltage-dependent K^+^ and Na^+^ currents that evoked action potentials. I-V relationship of Na^+^ currents from −60 to +60 mV **(A)** and K^+^ currents from −10 to +180 mV **(B)** in h DPSCs. Perfusion of hDPSCs with 1 μM tetrodotoxin (TTX) blocked Na^+^-inward currents. The insets display the original I-V traces obtained at different test potentials. **( C)** Representative traces from whole-cell voltage and current clamp recordings showing spontaneous activity in hDPSCs. **(D)** Single action potentials were elicited by depolarizing hDPSCs using three consecutive step current injections of 120 pA for 1500 ms separated by 1000 ms.

## Discussion

The inability of the mammalian CNS to spontaneously regenerate after damage [41] has led to the search for alternative non neural stem cell candidates for neuroregenerative cell therapies. Among them, mesenchymal stem cells (MSCs) have been postulated as a promising source [42]. MSCs can be extracted from different tissues such as the dental pulp. However, owing to their neural crest origin, the differentiation potential of hDPSCs is not restricted to mesenchymal cell lineages but can also extend to neural progenitors and mature neuronal markers expressing cells [13]. Nevertheless, the true capacity of hDPSCs to differentiate into functional neuron-like cells that can integrate into the CNS synaptic network, is still largely debated [10]. One important factor is the lack of consensus in hDPSCs differentiation protocols and neuronal differentiation criteria that may be found in the literature. In the present study, we presented a novel approach to generate electrophysiologically excitable neuron-like cells from hDPSCs by upgrading, via KCl pulses and RA, a standard protocol commonly used for NSC differentiation. Furthermore, we analyzed the reversibility of the changes induced by previous exposure to FBS during hDPSCs neural differentiation.

The use of FBS in the culture medium is a widespread practice in many laboratories, as it increases the growth rate and quality of the culture. However, different studies have reported the preference of hDPSCs to commit to mesenchymal-like phenotypes under fetal serum exposure [28,29]. The continuous presence of FBS in the medium leads to the formation of a plastic-adherent mesenchymal cell monolayer, which favors hDPSC differentiation into osteoblasts and odontoblasts [43]. Thus, in recent years, serum-free non-adherent sphere growth systems have been explored as more suitable techniques for hDPSCs expansion before their differentiation into other non-mesenchymal related lineages [13,17,44,45]. Neurosphere formation is a standard procedure used for brain NSC expansion because of the suitable microenvironment that is generated within the sphere. This intraspheral localization favors close physical contacts between neural progenitors, which are essential for mature neural cell commitment [30,31]. In our hands, we could also validate the importance of the sphere growth system for the maintenance of neural stemness. As we reported, hDPSCs grown as free-floating spheres presented higher basal expression levels of Nestin and GFAP, which are characteristic markers of NSCs.

In this work, we studied the reversibility of the changes induced by transient culture of hDPSCs in the presence of FBS with respect to neural differentiation. Thus, we compared two neural induction protocols. Sister cultures of hDPSCs from the same donors grown with FBS as a monolayer were compared with those grown as spheres in a serum-free environment after both were switched to the same neural induction medium for 21 days. Despite obtaining mature glial (S100β and p75^NTR^) and neuronal (DCX and NeuN) marker-expressing cells via both protocols, the percentage of positive cells for each marker and the morphology and functionality of the differentiated cells were very different between the two conditions, highlighting the importance of selecting an appropriate protocol for neural differentiation. Differentiated hDPSCs derived from serum-free dentospheres presented higher rates of S100β and p75^NTR^ positive cells, revealing a greater percentage of cells with a Schwann cell phenotype [35]. This finding is in line with the fact that neural crest (NCs) progenitors, which are present in dental pulp tissue, are committed differently in the presence or absence of FBS. In addition, as reported previously by other authors, hDPSCs that had been previously grown as floating spheres presented, during the neural differentiation process, morphological adaptations that resembled those of neural cells, with a rounded large cytoplasm surrounded by a peripheral halo with very long and thin cytoplasmic extensions [17,19]. This phenotype could not be observed in hDPSCs previously grown in the presence of FBS, which presented morphological characteristics of mesenchymal-like cells, like large lamellipodia, even after the differentiation process.

To define an induced cell as a true neuron, at least three attributes should be accomplished: showing a characteristic neuronal morphology, expressing neuron-specific gene products, and displaying electrophysiological activities such as synaptic transmission and action potential firing [46]. Even if hDPSCs grown as an adherent monolayer expressed mature neuronal markers after they were switched into neural differentiation media, neither mature neuronal morphology nor electrophysiological functionality could be induced in them. Similar results were reported by Li et al., who, after comparing three neurogenic differentiation protocols, only generated excitable cells performing one single AP with no baseline potential recovery through a neurosphere-mediated approach in hDPSCs [19]. Similarly, we could not generate any AP in hDPSCs when these were previously grown in FBS-containing medium, and only a few voltage-gated sodium and potassium currents could be recorded in those cells previously grown as floating spheres after 21 days of exposure to conventional neural differentiation protocols.

Signals present in neural induction media, such as KCl and RA, are known to promote neurogenesis and GABAergic phenotypes in NSCs [23]. RA has been shown to support the survival of neural progenitors [47] and induce neuronal differentiation in stem cells [48]. In the adult brain, RA influences synaptic plasticity in the hippocampus [49] and regulates neurogenesis in the subventricular zone and the hippocampal subgranular area [50]. On the other hand, increased electrical activity triggered by KCl has been reported to downregulate Nestin levels, increase the number of postmitotic neurons, and induce neurite outgrowth [23]. Thus, we decided to improve the neuronal differentiation protocol with KCl pulses and RA, expecting to increase the neuronal fate commitment of differentiated DPSCs. The differentiation times were extended to two months, with the goal of achieving better electrophysiological recordings.

After the neural induction process, mature neuronal markers such as NeuN or MAP2 [51] were expressed by differentiated hDPSCs. However, non-statistically significant differences were shown between the improved and non-improved culture conditions, which could be related to some variability in the cell responses observed between different donors, as previously reported by other authors [17,19]. In addition, we also assessed the expression of synaptic proteins in differentiated hDPSCs at the protein and mRNA levels. Synapsin-I is a phosphoprotein associated with synaptic vesicles (SVs) that serves as a linker between SVs and actin filaments in presynaptic terminals to cluster SVs in a phosphorylation-dependent manner during neurotransmitter release [52]. In our cultures, this presynaptic protein was expressed, indicating the possibility that our differentiated hDPSCs produce synaptic vesicles. However, for complete synaptic transmission, many other key components, such as postsynaptic proteins, neurotransmitters (NTs) and their receptors, are also needed.

In the mature CNS the majority of synapses are chemical, releasing a specific NT into the synaptic cleft. This NT subsequently binds to specific receptors in the postsynaptic terminal, and an ion flow through these receptors depolarizes or hyperpolarizes the postsynaptic neuron. Most excitatory synapses in the CNS are glutamatergic, unlike inhibitory synapses, which are predominantly GABAergic [53]. After hDPSC neurodifferentiation we obtained a heterogeneous cell population in which the cells expressed the basic building blocks for both types of synapses. For example, in our cultures, we identified the central postsynaptic organizers of glutamatergic synapses, PSD95 [37], the vesicular glutamate transporter VGLUT-2, and the ionotropic glutamate receptor kainate subunit GRIK2. Furthermore, many differentiated hDPSCs also presented a strong GABAergic phenotype after exposure to KCl and RA, as also reported in NSCs by Bosch et al. [23]. This was assessed by their capacity to produce the enzyme responsible for GABA synthesis (GAD 65) and GABA itself. In addition, postsynaptic GABA receptor subunits (GABRB1) and the central GABAergic postsynaptic organizer Gephyrin were also expressed.

As previously mentioned, for reasonably claiming an induced cell to be a fully mature neuron, its functionality must be demonstrated. For that, electrophysiological experiments must be carried out. Action potentials are the fundamental electrical signals of the CNS used for information coding and transmission and are shaped and generated in the AIS, a neuronal region with a high density of membrane voltage-gated Na_v_ channels [38]. The anchoring of Na_v_ channels to the AIS is known to be mediated by the protein Ankyrin-G [40] whose expression was detected, to our knowledge for the first time, in our hDPSCs cultures after the upgraded protocol of neural commitment. Taken together with the greater prevalence of sodium and potassium voltage-gated currents observed in our KCl- and RA-treated cells, we hypothesize that our differentiated cells possess most of the machinery required for action potential generation and propagation. Nevertheless, considering the results previously obtained by other authors [17,19], we decided to extend the maturation period for up to two months. After this time, the electrophysiological recordings of hDPSCs differentiated with KCl and RA revealed that they were able to generate voltage-gated sodium and potassium currents. The recorded I-V curves indicate that the opening kinetics of these channels are consistent with those of the neuronal Na_v_ and K_v_ channels. As also described by other authors the neuronal specificity of those sodium currents was confirmed by selectively blocking them with TTX [17–19]. Finally, differentiated hDPSCs clearly exhibited spontaneous electrophysiological activity in the absence of stimulation, as recorded in both voltage and current clamp mode. Electrical activity is a very important requisite for the establishment of synaptic contacts between neuronal cells, where an immature pattern of activity precedes the emergence of chemical synapses and integrative neuronal functions [54,55]. Our study constitutes a first demonstration of functional and spontaneous electrophysiological activity in neurodifferentiated hDPSCs, which sheds light on the capacity of hDPSCs to generate repetitive AP firing patterns characteristic of fully mature neuronal cells and to maintain an electrical excitability in the long term. We are aware that the next step is to assess whether these electrophysiologically immature neuron-like cells differentiated from hDPSCs might be able to contribute to the formation of and/or integrate into preexisting synaptic networks *in vivo*, but it is beyond the scope of the present manuscript. However, this study represents a substantial step ahead in the generation of truly excitable neuron-like cells, which may hold promise in the treatment of brain lesions and neurodegenerative disorders.

### Conclusions

We found that FBS-induced changes influence the long-term neurodifferentiation fate of hDPSCs and that *in vitro* culture media environment cues are decisive in the effectiveness of hDPSCs neural induction. Neurodifferentiated hDPSCs generated through serum-free spheroid expansion were quite different to those cells obtained after expansion in the presence of FBS, regarding not only the ability to give rise to S100β^+^/p75^NTR+^ Schwann cells, but also exhibiting a drastically different cellular morphology, with poly-dendritic cell shapes and marked increases in the branch length, perimeter and area. The addition of RA and KCl increased the expression of GABA, kainate receptor subunits (GRIK2), voltage-gated sodium (SCN8A), and potassium (KCNA2) channels after differentiation, resulting in the generation of electrophysiologically excitable cells which triggered repetitive action potentials. This study showcases the potential of an easily accessible human stem cell source for nerve tissue engineering. Further studies on differentiated neuronal-like hDPSCs brain grafts should focus on the potential of these cells for engraftment and integration into preexistent brain synaptic circuits to assess their potential utility for AD cell therapy.

## Supporting information

Supplementary figurexs, tables and movies

## Abbreviations

aCSF: Artificial cerebrospinal fluid
AIS: Axon initial segment
*ANK3*: Ankyrin-G gene
APs: Action potentials
BDNF: Brain derived neurotrophic factor
BSA: Bovine serum albumin
CaCl_2_: Calcium chloride
CaCl_2_-H_2_O: Calcium chloride hydrate
CNS: Central nervous system
DCX: Doublecortin
DIV: Days *in vitro*
DMEM: Dulbecco’s Modified Eagle Medium
FBS: Fetal bovine serum
GABA: Gamma Aminobutyric acid
GABRB1: Gamma-aminobutyric acid type A receptor subunit
GAD65: Glutamic acid decarboxylase 65
GAPDH: Glyceraldehyde 3-phosphate dehydrogenase
GFAP: Glial fibrillary acidic protein
GPHN: Gephyrin
GRIK2: Glutamate ionotropic receptor kainate type subunit 2
hDPSCs: Human dental pulp stem cells
HEPES: 4-(2-hydroxyethyl)-1-piperazineethanesulfonic acid
K^+^: Potassium ion
KCl: Potassium chloride
*KCNA2:*: Potassium voltage-gated channel subfamily A member 2 gene
KH_2_PO_4_: Monobasic potassium phosphate
KOH: Potassium hydroxide
K_v_: Voltage-dependent potassium channels
MAP2: Microtubule associated protein 2
MgCl: Magnesium chloride
MgSO4: magnesium sulphate
mRNA: Messenger ribonucleic acid
ms: Milliseconds
MSCs: Mesenchymal stem cells
mV: Milivolts
Na^+^: Sodium ion
Na_2_ATP: Adenosine 5′-triphosphate disodium salt hydrate
NaCl: Sodium chloride
NaHCO_3_: Sodium hydrogencarbonate
NaOH: Sodium hydroxide
Na_v_: Voltage-gated sodium channels
NCs: Neural crest progenitors
NeuN: Hexaribonucleotide Binding Protein-3
NeuN: Neuronal nuclear antigen
NGFR: Nerve growth factor receptor
NSCs: Neural stem cells
NT-3: Neurotrophin-3
NTs: Neurotransmitters
P75^NTR^: p75 neurotrophin receptor
pA/pF: Picoampere/Picofarad
pA: Picoampere
PBS: Phosphate buffered saline
PFA: Paraformaldehyde
PSD95: Postsynaptic density protein 95
qPCR: Quantitative polymerase chain reaction
qRT‒PCR: Real-time quantitative reverse transcription ‒ polymerase chain reaction
RA: Retinoic acid
S100β: S100 calcium-binding protein B
*SCN8A*: Sodium voltage-gated channel alpha subunit 8 gene
SVs: Synaptic vesicles
TTX: Tetrodotoxin
VGlut2: Vesicular glutamate transporter 2

## Acknowledgement

We would like to thank to Ricardo Andrade and Alex Díez from the Analytical and High Resolution Microscopy Service in Biomedicine of the SGIker services (UPV/EHU) and Rafael Martínez Conde’s maxillofacial surgery clinic. The authors declare that they have not use AI-generated work in this manuscript.

## Author contributions

B.P.R., F.U., G.I. and J.R.P. were responsible for the study concept and design. B.P.R, A.M.B, I.M.R., J.L., J.S.M., Y. P., and R.B.T. performed the investigation and formal analysis. B.P.R, G.I. and J.R.P. contributed to the methodology, and writing of the original draft. S.M., F.U., G.I. and J.R.P. handled conceptualization, funding acquisition and supervision. All authors reviewed and critically revised the draft manuscript. Authors approved the final manuscript.

## Data availability

All additional files are included in the manuscript.

## Declarations

### Ethics approval and consent to participate

Human third molars were obtained from healthy donors of between 19 and 45 years of age and tooth samples were obtained by donation after written informed consent in compliance with the 14/2007 Spanish directive for Biomedical research. The study protocol was approved on date 03/02/2021 by the Ethics Committee of the University of the Basque Country (UPV/EHU) and the competent authority (Administración Foral de Bizkaia) regarding the use of human cells with CEISH M10/2020/172 and M10/2023/025 entitled “*Matrices de anclaje nanoestructuradas basadas en grafeno y polímeros biodegradables para inducir la neurodiferenciación de células madre y regenerar el tejido nervioso*”. The study was conducted in accordance with the Declaration of Helsinki.

## Consent for publication

Not applicable.

