## Supplementary figurexs, tables and movies for "Functional differentiation of Human Dental Pulp Stem Cells into neuron-like cells exhibiting electrophysiological activity": Table 1.docx

**Table 1:** Primer pairs sequences used in this study.

| **Primers** |  | **5’-3’ Sequence** | **Annealing (ºC)** |
| --- | --- | --- | --- |
| **GAPDH** | Forward | CTTTTGCGTCGCCAG | 60.3 |
|  | Reverse | CACCTATAACAACGGTAGTT | 60.8 |
| **ACTB**  **(β-ACTIN)** | Forward | GACGACATGGAGAAAATCTG | 59.7 |
|  | Reverse | CTCTTCTACTGGGTCTAGTA | 58 |
| **SYN-I** | Forward | TTCAACCTTCCAGAGCCAGCC | 69.7 |
|  | Reverse | AGAACCGGGAGATGGGTTCTC | 67.5 |
| **PSD95** | Forward | CGCCTGCATCTCCGAATCT | 67.7 |
|  | Reverse | CGGGCGGGATTAAGGAGTTT | 67.7 |
| **NES** | Forward | TTTCAGGACCCCAAGC | 59.7 |
|  | Reverse | CTGGAGGAATTCTTGGTTC | 58.7 |
| **GFAP** | Forward | CACCACGATGTTCCTCTTGA | 57.3 |
|  | Reverse | ATCGAGATCGCCACCTACAG | 59.4 |
| **S100β** | Forward | ACCAATATTCTGGAAGGGAG | 59.4 |
|  | Reverse | CCTCTAAGAAATGGGAAAGC | 59.3 |
| **NGFR** | Forward | TGAGGCACCTCCAGAACAAG | 65.4 |
|  | Reverse | GCAGCTGTTCCACCTCTTGAA | 66.7 |
| **DCX** | Forward | TGCCTCAGGGAGTGCGTTA | 58.8 |
|  | Reverse | GAACAGACATAGCTTTCCCCTTC | 60.6 |
| **RBFOX3 (NEUN)** | Forward | CAGAGTGCCTCGGGGAAAAT | 60.0 |
|  | Reverse | CACCAGGTTCCGAGTCCAAA | 59.9 |
| **MAP2** | Forward | TGTGTCGTGTTCTCAAAGGG | 57.3 |
|  | Reverse | TGCATATGCGCTGATTCTTC | 55.3 |
| **SCN8A** | Forward | CATGTATGCAGCTGTAGATTC | 57.4 |
|  | Reverse | CCGAAGATGATGAAGATGAC | 59.7 |
| **KCNA2** | Forward | TTGCATAATAGCCTGCATTG | 61.2 |
|  | Reverse | TTCACTTGCACAACTATTGC | 58.9 |
| **GRIK2** | Forward | TTTTGGAAGAGCCTTATGTC | 58.4 |
|  | Reverse | CTGAGGAGATCAATGCAATAG | 58.9 |
| **GABRB1** | Forward | GCTACAAGAAGATTGGCTAC | 55.7 |
|  | Reverse | CCCTCCAATCCATTTATCTG | 60.7 |
| **VGLUT2** | Forward | TCAGATTCCGGGAGGCTACA | 66.9 |
|  | Reverse | TGGGTAGGTCACACCCTCAA | 65.4 |
| **GPHN** | Forward | GTCACTCCAGAGGCCACAAA | 65.3 |
|  | Reverse | TAGAGAGCATGCCCAGAGGT | 63.9 |
| **ANK3** | Forward | CTGCTAGAGGCTTGCCAGAG | 64.3 |
|  | Reverse | TGACCACTTCGCAGACATCC | 66.4 |
