## Supplementary figurexs, tables and movies for "Functional differentiation of Human Dental Pulp Stem Cells into neuron-like cells exhibiting electrophysiological activity": Table 2.docx

|  | Neurocult-Neural diff | | | | | | Neurocult-Neural diff KCl | | | | | |
| --- | --- | --- | --- | --- | --- | --- | --- | --- | --- | --- | --- | --- |
|  | I_Na_ | | **I_K_** | | **AP** | | **I_Na_** | | **I_K_** | | **AP** | |
| DIV | Trial | Resp. | Trial | Resp. | Trial | Resp. | Trial | Resp. | Trial | Resp. | Trial | Resp. |
| 21 | 5 | 0 | 3 | 1 | 3 | 0 | 20 | 3 | 13 | 3 | 15 | 0 |
| 60 |  | | | | | | 24 | 5 | 21 | 3 | 29 | 6 |

**Table 2:** Electrophysiological recordings.
