## Supplementary figures and images for "Functional differentiation of Human Dental Pulp Stem Cells into neuron-like cells exhibiting electrophysiological activity"

### Supplemental Figure1.tif

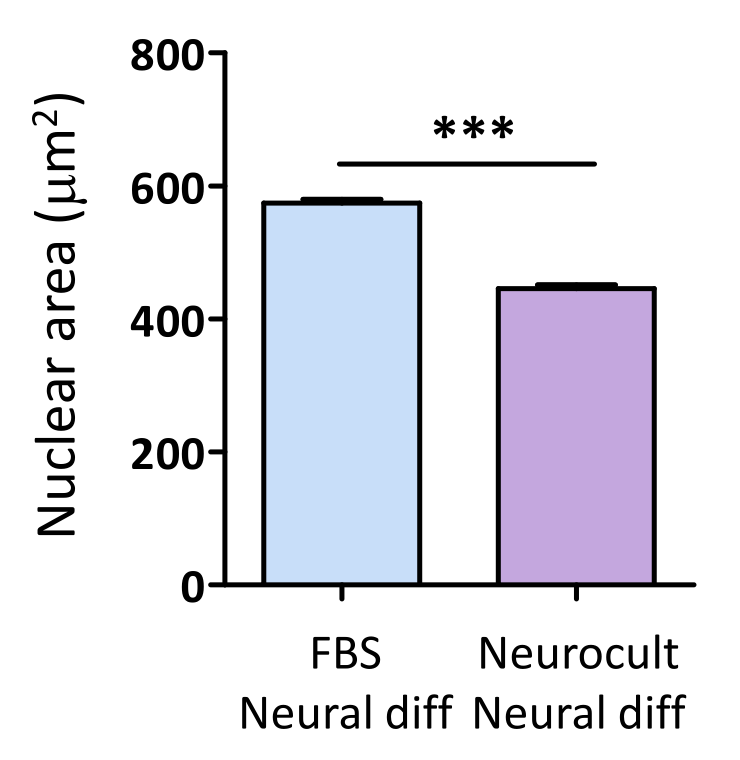

### Supplemental Figure2.tif

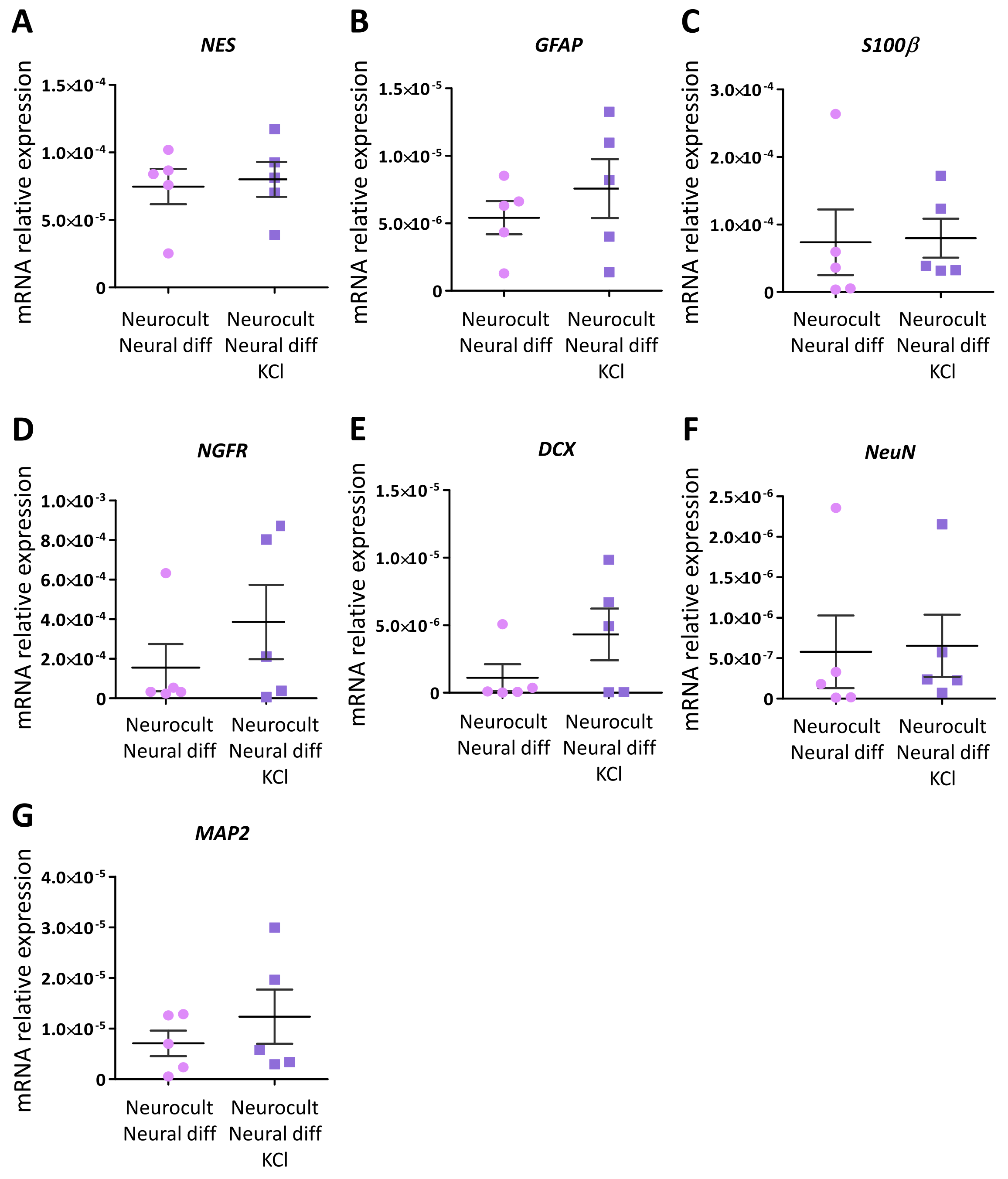
